# Cryo-EM structures of PP2A:B55-FAM122A and PP2A:B55-ARPP19

**DOI:** 10.1101/2023.08.31.555365

**Authors:** Sathish K.R. Padi, Margaret R. Vos, Rachel J. Godek, James R. Fuller, Thomas Kruse, Jamin B. Hein, Jakob Nilsson, Matthew S. Kelker, Rebecca Page, Wolfgang Peti

## Abstract

Progression through the cell cycle is controlled by regulated and abrupt changes in phosphorylation.^1^ Mitotic entry is initiated by increased phosphorylation of mitotic proteins, a process driven by kinases,^2^ while mitotic exit is achieved by counteracting dephosphorylation, a process driven by phosphatases, especially PP2A:B55.^3^ While the role of kinases in mitotic entry is well-established, recent data have shown that mitosis is only successfully initiated when the counterbalancing phosphatases are also inhibited.^4^ For PP2A:B55, inhibition is achieved by the two intrinsically disordered proteins (IDPs), ARPP19 (phosphorylation-dependent)^6,7^ and FAM122A^5^ (inhibition is phosphorylation-independent). Despite their critical roles in mitosis, the mechanisms by which they achieve PP2A:B55 inhibition is unknown. Here, we report the cryo-electron microscopy structures of PP2A:B55 bound to phosphorylated ARPP19 and FAM122A. Consistent with our complementary NMR spectroscopy studies both IDPs bind PP2A:B55, but do so in highly distinct manners, unexpectedly leveraging multiple distinct binding sites on B55. Our extensive structural, biophysical and biochemical data explain how substrates and inhibitors are recruited to PP2A:B55 and provides a molecular roadmap for the development of therapeutic interventions for PP2A:B55 related diseases.

## Introduction

An essential function of the PP2A:B55 holoenzyme is the control of cell cycle progression through mitosis.^4,6–9^ Entry into mitosis is accomplished via the activation of the CDK1/CyclinB kinase, whose activity allows for nuclear envelope breakdown, chromatin condensation and spindle formation.^10,11^ Exit from mitosis is initiated by CyclinB ubiquitination via the anaphase promoting complex (APC) leading to its degradation.^12^ Simultaneously, dephosphorylation of mitotic substrates by Protein Phosphatases (PPPs), especially PP2A:B55, is initiated.^13^ Indeed, like CyclinB degradation during mitotic exit, inhibition of PP2A:B55 activity during mitotic onset has been shown to create the necessary dynamic feedback for mitotic substrate phosphorylation.^14^

Recently, the PP2A:B55 holoenzyme was shown to also regulate the entry into mitosis at the G2/M checkpoint, as inhibition of PP2A:B55 permits normal progression through the checkpoint.^8,9^ These essential PP2A:B55 inhibitory events are achieved by its interaction with two distinct intrinsically disordered protein (IDP) inhibitors, ARPP19 and FAM122A.^7,15–17^ The mechanisms by which these inhibitors block PP2A:B55 activity differ, as ARPP19 strictly requires phosphorylation by MASTL kinase to inhibit PP2A:B55^6,7^ while FAM122A inhibits PP2A:B55 in a phosphorylation-independent manner.^5,8^ The current data suggest that these IDP inhibitors engage PP2A:B55 sequentially during mitotic entry (**Fig. 1a**). Namely, PP2A:B55 is initially bound and inhibited by FAM122A. This inhibition results in the full activation of mitotic kinases, including MASTL, which phosphorylates ARPP19 on Ser62 (*pS62*ARPP19). Via a currently unknown mechanism, *pS62*ARPP19 displaces FAM122A from PP2A:B55. *pS62*ARPP19 functions first as an inhibitor of PP2A:B55 but later becomes a substrate.^18^ This dephosphorylation reactivates PP2A:B55 and allows for the progression through mitotic exit. Despite our understanding of the importance of PP2A:B55 in mitosis, and the identity of the IDP inhibitors that mediate PP2A:B55 inhibition during mitotic entry, a detailed understanding of how this inhibition is achieved at a molecular level has remained unknown.

**Figure 1.**
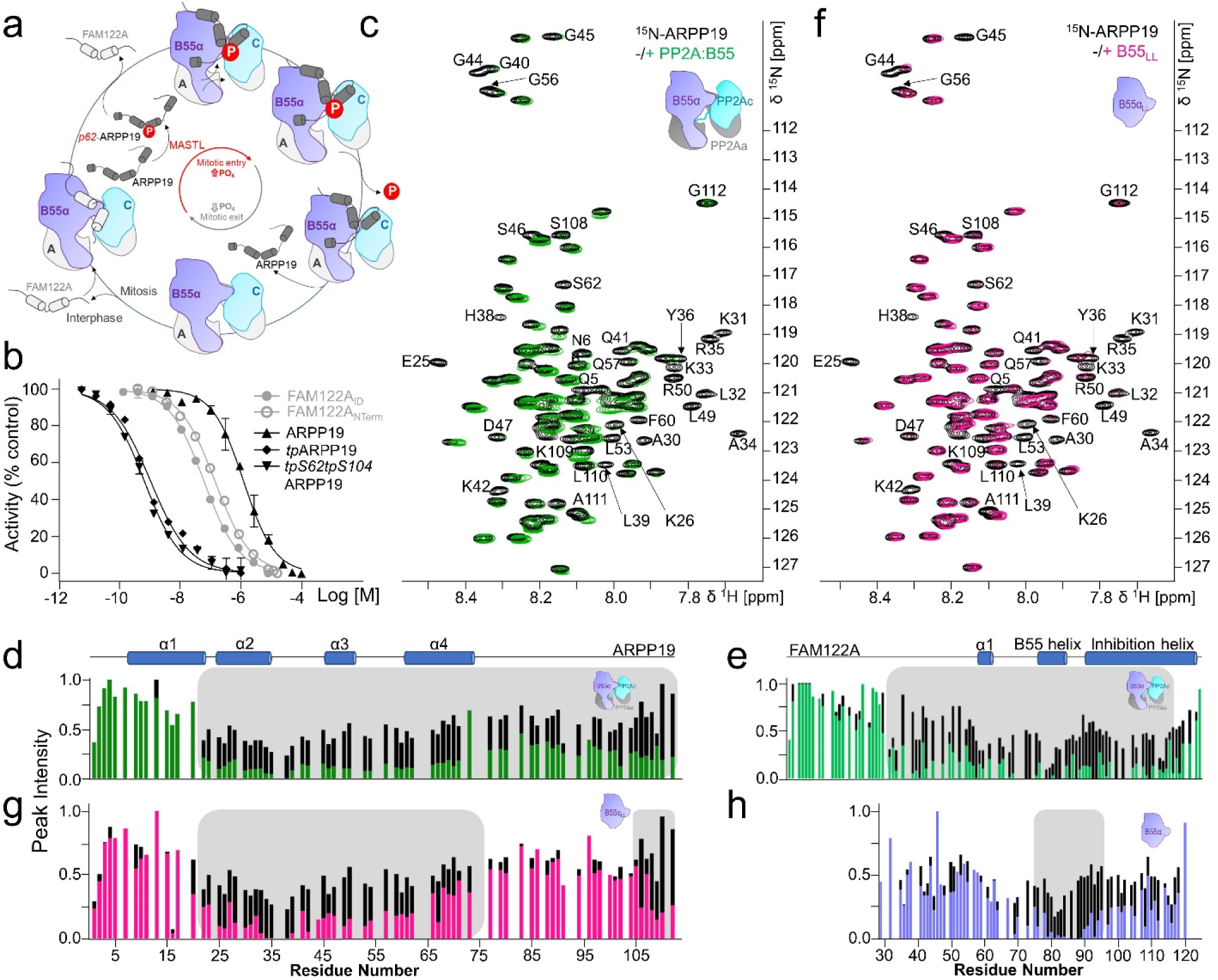
ARPP19 and FAM122A inhibit PP2A:B55. **a**. Current understanding of the roles of ARPP19 and FAM122A in regulating PP2A:B55 activity during the cell cycle. **b**. IC_50_ curves for PP2A:B55 inhibition by ARPP19 (with and without phosphorylation), FAM122A_Nterm_ and FAM122A_ID_. IC_50_ values are reported in **Table 1**. Data are presented as mean values ± SD, n = 3 experimental replicates. **c**. 2D [^1^H,^15^N] HSQC spectrum of ^15^N-labeled ARPP19 alone (black) and in complex with PP2A:B55 (green). **d**. Peak intensity vs ARPP19 protein sequence plot for ARPP19 alone (black) and when bound to PP2A:B55 (green); grey box highlights ARPP19 residues with reduced intensities when bound to PP2A:B55. Secondary structure elements based on NMR CSI data are indicated. **e**. Peak intensity vs FAM122A protein sequence plot for FAM122A_Nterm_ alone (black) and when bound to PP2A:B55 (green); grey box highlights FAM122A residues with reduced intensities when bound to PP2A:B55. Secondary structure elements based on NMR CSI data are indicated. **f**. 2D [^1^H,^15^N] HSQC spectrum of ^15^N-labeled ARPP19 alone (black) and in complex with B55_LL_ (pink). **g**. Peak intensity vs ARPP19 protein sequence plot for ARPP19 alone (black) and when bound to B55_LL_ (pink); grey box highlights ARPP19 residues with reduced intensities when bound to B55_LL_. **h**. Peak intensity vs FAM122A_ID_ protein sequence plot for FAM122A_ID_ alone (black) and when bound to B55_LL_ (blue); grey box highlights FAM122A_ID_ residues with reduced intensities when bound to B55_LL_.

**Table 1:** Inhibition (IC_50_) of PP2A:B55 by ARPP19 and FAM122A: All experiments were performed as experimental triplicates; mean ± std; n = number of independent measurements; #indicates clinical cancer variant.

| Inhibitor | IC<sub>50</sub> [nM] | n |
| --- | --- | --- |
| ARPP19 | 1576 ± 151 | 4 |
| *tp*ARPP19 | 1.3 ± 0.3 | 4 |
| *tpS62tpS104*ARPP19 | 0.6 ± 0.1 | 3 |
| ARPP19<sub>19-75</sub> | ND | 3 |
| *tp*ARPP19<sub>19-75</sub> | 71 ± 9.5 | 3 |
| FAM122A<sub>Nterm</sub> | 151 ± 7 | 3 |
| FAM122A<sub>ID</sub> (S120C) | 57 ± 4 | 3 |
| FAM122A<sub>ID</sub> (S120C)_R84A/L85A | 2147 ± 208 | 3 |
| FAM122A<sub>ID</sub> (S120C)_I88A/K89A | 2740 ± 90 | 3 |
| FAM122A<sub>ID</sub> (S120C)_E91K | 1037 ± 29 | 3 |
| FAM122A<sub>ID</sub>_(S120C)_E92K | 579 ± 2 | 3 |
| FAM122A<sub>ID</sub> (S120C)_E104A | 108 ± 4 | 3 |
| FAM122A<sub>ID</sub> (S120C)_E106A | 38 ± 3 | 3 |
| FAM122A<sub>ID</sub>_E92K<sup>#</sup> | 540 ± 42 | 3 |
| FAM122A<sub>ID</sub>_R105L<sup>#</sup> | 616 ± 35 | 3 |
| FAM122A<sub>ID</sub>_V107G<sup>#</sup> | 342 ± 13 | 3 |

### The interaction of ARPP19 and FAM122A with PP2A:B55

To determine how ARPP19 and FAM122A bind PP2A:B55, we established a method for producing high yields of active PP2A:B55 from Expi293F cells (**Extended Data Fig. 1a-h**; the C-terminal residue of PP2Ac is fully methylated^19–21^, *m*L309). We quantified PP2A:B55 inhibition by ARPP19, thiophosphorylated ARPP19 (full-length, aa 1-112, phosphorylated with ATP-γ-S using MASTL kinase) and FAM122A (N-terminal domain, aa 1-124; FAM122A_Nterm_) (**Fig. 1b**). While PP2A:B55 was only moderately inhibited by ARPP19, it was strongly inhibited by both thiophosphorylated ARPP19 and FAM122A_Nterm_ (**Table 1; Extended Data Fig. 2**), with thiophosphorylated ARPP19 inhibiting PP2A:B55 ∼250-fold more potently than FAM122A. Fluorescent polarization binding measurements also showed that both FAM122A and ARPP19 bind PP2:B55 tightly, with thiophosphorylation not influencing binding (**Table 2; Extended Data Fig. 3**). We then used NMR spectroscopy to identify the residues in ARPP19 and FAM122A that interact with PP2A:B55. The 2D [^1^H,^15^N] heteronuclear single quantum coherence (HSQC) spectra of unbound ARPP19^22,23^ and FAM122A_Nterm_ confirmed that both are IDPs with multiple regions of amino acids with preferred α-helical propensities (chemical shift index, CSI, analysis; **Extended Data Fig. 4**a-f). Overlaying the 2D [^1^H,^15^N] HSQC spectra of ARPP19 and FAM122A in the presence and absence of PP2A:B55 identified the residues that bind PP2A:B55 (peaks with reduced intensities are the result of either a direct interaction, a dynamic charge:charge interaction or intermediate timescale conformational exchange) (**Figs. 1c-e**). For ARPP19, ∼90 N/H^N^ cross peaks (residues 20-112) showed reduced intensities. Since ARPP19 inhibition of PP2A-B55 strictly requires phosphorylation, we also thiophosphorylated ARPP19 using MASTL kinase. The 2D [^1^H,^15^N] HSQC spectrum of thiophosphorylated ARPP19 identified two phosphorylated residues, Ser62, the established MASTL phosphorylation substrate, and, Ser104, a serine that was previously identified as a PKA substrate, but also shows recognition site homology to the MASTL specificity sequence^24^. The NMR data show that MASTL phosphorylation does not alter the preferred ensemble of structures of ARPP19 (**Extended Data Fig. 4**g-i). Thus, we generated ARPP19 S104A and repeated the thiophosphorylation to obtain singly phosphorylated ARPP19 (*tp*S62ARPP19_S104A;_ hereafter, *tp*ARPP19). Overlaying the 2D [^1^H,^15^N] HSQC spectra of *tp*ARPP19 and *tpS62tpS104*ARPP19 with PP2A:B55 showed that the intensities of the same ∼90 N/H^N^ cross peaks identified with unphosphorylated ARPP19 are reduced in the presence of PP2A:B55 (**Extended Data Fig. 5**a-d). Similar NMR interaction experiments with FAM122A_Nterm_ showed that the intensities of ∼85 cross peaks (residues 30-115) were reduced in the presence of PP2A:B55 (**Fig. 1e; Extended Data Fig. 5**e). Based on this result, we created FAM122A_ID_ (aa 29-120) that includes all PP2A:B55 interacting residues (**Extended Data Figs. 3,5f,g**).

**Table 2:** Binding affinity (K_D_) of ARPP19 and FAM122A to PP2A:B55 or B55_LL_: All experiments were performed as experimental triplicates; mean ± std; n = number of independent measurements.

| Inhibitor | B55 | $K_D$ [nM] | n |
| --- | --- | --- | --- |
| ARPP19 (S10C) | PP2A:B55 | 113 ± 6 | 4 |
| ARPP19 S104A (S10C) | PP2A:B55 | 115 ± 5 | 4 |
| *tp*ARPP19 (S10C) | PP2A:B55 | 113 ± 12 | 4 |
| *tpS62tpS104*ARPP19 (S10C) | PP2A:B55 | 74 ± 1 | 4 |
| ARPP19 (S10C) | B55<sub>LL</sub> | 323 ± 41 | 4 |
| ARPP19 S104A (S10C) | B55<sub>LL</sub> | 326 ± 40 | 3 |
| FAM122A<sub>ID</sub> (S120C) | PP2A:B55 | 190 ± 12 | 3 |
| FAM122A<sub>67-120</sub> (S120C) | PP2A:B55 | 518 ± 25 | 3 |
| FAM122A<sub>ID</sub> (S120C) _R84A/L85A | PP2A:B55 | 307 ± 3 | 3 |
| FAM122A<sub>ID</sub> (S120C) _I88A/K89A | PP2A:B55 | 386 ± 5 | 3 |
| FAM122A<sub>ID</sub> (S120C) _E91K | PP2A:B55 | 317 ± 7 | 3 |
| FAM122A<sub>ID</sub> (S120C) _E92K | PP2A:B55 | 356 ± 13 | 3 |
| FAM122A<sub>67-120</sub> (S120C) | B55<sub>LL</sub> | 827 ± 44 | 3 |
| FAM122A<sub>ID</sub> (S120C) | B55<sub>LL</sub> | 837 ± 53 | 3 |

ARPP19 and FAM122A inhibit only B55-containing PP2A holoenzymes.^5,8^ To identify which residues of ARPP19 and FAM122A bind B55, we repeated the NMR experiments using B55 loopless (B55_LL_), a B55 variant that lacks its PP2Aa interaction loop (aa 126-164 replaced with residues ‘NG’; **Extended Data Fig. 1**a), and thus is unable to bind PP2Aa/PP2Ac. Overlaying the 2D [^1^H,^15^N] HSQC spectra of ARPP19 and *tpS62tpS104*ARPP19 with and without B55_LL_ showed that the identity and number of N/H^N^ cross-peaks with reduced intensities are similar, but not identical, to those observed in the PP2A:B55 experiments (**Figs. 1f,g; Extended Data Figs. 6a,b**). Namely, while the peaks corresponding to residues 20-75 and 105-112 show significant reductions in intensities, ARPP19 residues 75-104 show little/no intensity loss with B55_LL_. This shows that two distinct ARPP19 domains, 20-75 and 105-112, bind B55. An overlay of the 2D [^1^H,^15^N] HSQC spectrum of FAM122A_ID_ with and without B55_LL_ (**Fig. 1h, Extended Data Fig. 6**c) showed that N/H^N^ cross-peaks with reduced intensities correspond to FAM122A residues 73-95, which bind solely to B55. Both ARPP19 and FAM122A bind B55_LL_ with reduced affinities compared to PP2A:B55 (ARPP19, 2.8-fold; FAM122A, 4.4-fold; **Table 2, Extended Data Fig. 3**). The length of these B55 interaction regions, more than 20 residues, was longer than expected. This is because most PPPs, including PP2A:B56, bind their substrates and regulators using <u>S</u>hort <u>Li</u>near <u>M</u>otifs (SLiMs) that are typically 4-8 residues long, as they bind their cognate PPP in an extended fashion.^25–29^ This suggests that the mechanism of ARPP19 and FAM122A binding to B55 differs from that of canonical members of the PPP family. Consistent with this, our NMR data showed that these IDP inhibitors exhibit significant helical propensities in solution (**Extended Data Figs. 4c,f**), suggesting that they likely bind B55 as helices.

### Cryo-EM structures of PP2A:B55-tpARPP19 and PP2A:B55-FAM122A_ID_

Following extensive sample and data collection optimization, we determined the structures of PP2A:B55-*tp*ARPP19 and PP2A:B55-FAM122A_ID_ using single particle cryogenic electron microscopy (cryo-EM) at average resolutions of 2.77 and 2.80 Å, respectively (**Figs. 2a,b; Extended Data Figs. 7-10**). Modeling of the PP2Aa, B55 and PP2Ac subunits was based on the previously solved PP2A:B55 crystal structure.^30^ Compared to the PP2A:B55 crystal structure, the horseshoe-shaped conformation of PP2Aa contracted upon inhibitor binding (**Fig. 2c**), allowing B55 loop ^84^FDYLKS^89^ to bind the PP2Ac hydrophobic groove. Despite this contraction, the interfaces between PP2Ac and PP2Aa (repeats 11-15) and B55 and PP2Aa (repeats 1-7) were identical to those observed in the crystal structure.

**Figure 2.**
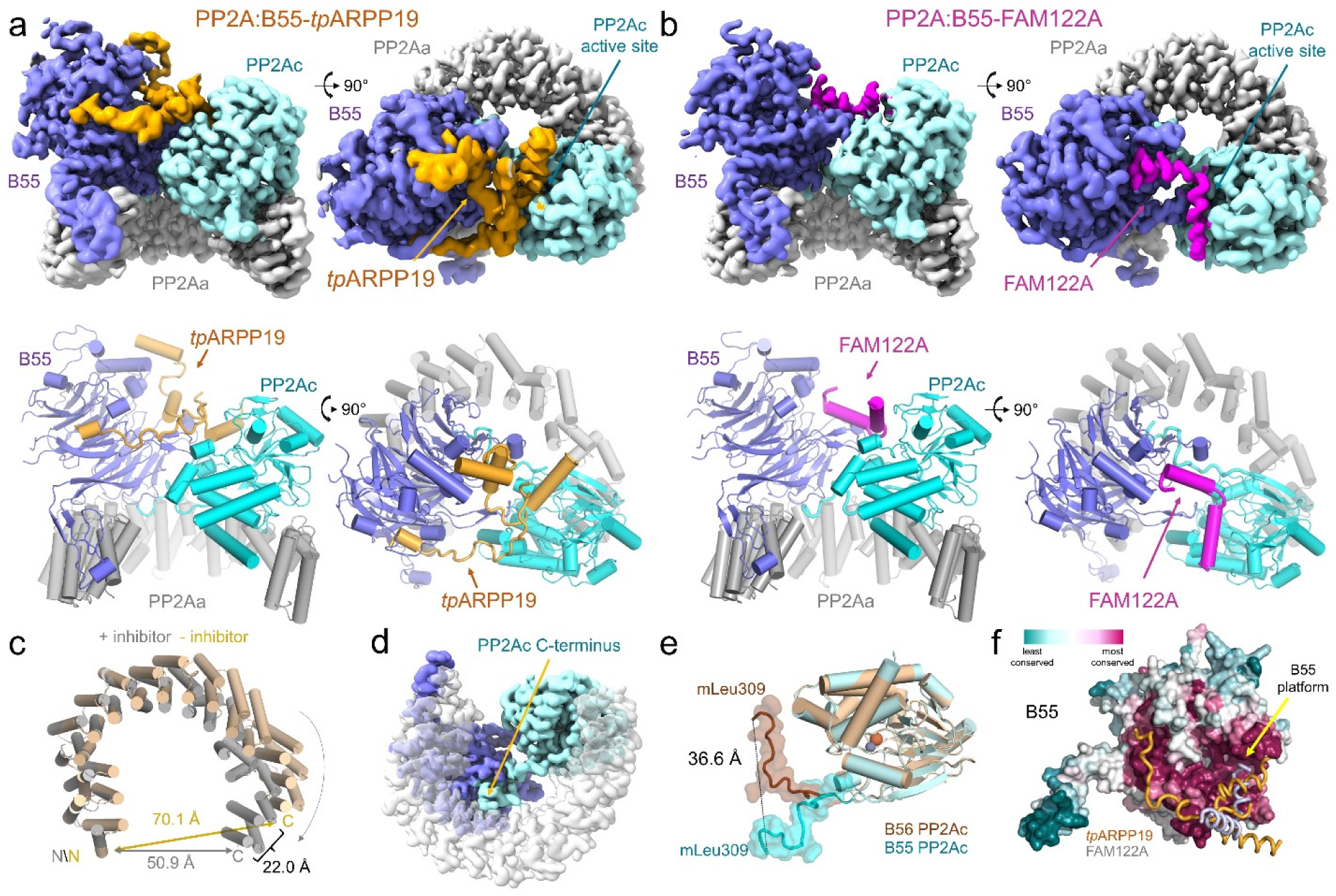
Structure of the PP2A:B55-*tp*ARPP19 and PP2A:B55-FAM122A complex. **a.** Cryo-EM map and model of the PP2A:B55-*tp*ARPP19 complex. Two views of the map (top) are shown next to the corresponding view of the molecular model (bottom). **b**. Cryo-EM map and model of the PP2A:B55-FAM122A complex. Same views as in a. **c**. Overlay of PP2Aa from the PP2A:B55 crystal structure (pdbid 3DW8; beige) and the PP2A:B55-inhibitor model (grey), superimposed using heat repeats 1-6. **d**. Rotated view of the experimental map; PP2Aa is transparent to show the extended PP2Ac C-terminal tail. **e**. Overlay of the PP2Ac catalytic domains from PP2A:B55-FAM122A (cyan) and PP2A:B56 (brown; pdbid 2IAE), with the C-terminal tails (aa R294-mL309) shown as a surface. The presence of B55 or B56 in the PP2A holoenzyme results in distinct binding sites for the PP2Ac C-terminal tail. **f**. B55 shown as a surface and colored by sequence conservation using the scale shown; ARPP19 (orange) and FAM122A (grey) bind to the most conserved B55 surface. The location of the B55 platform is also indicated.

In both PP2A:B55-inhibitor maps, continuous sections of density not accounted for by the PP2A:B55 crystal structure were observed. The density common to both maps belongs to the PP2Ac C-terminus (**Fig. 2d**, aa 294-309), which was not modeled in the PP2A:B55 crystal structure.^30^ The C-terminus extends across the PP2Aa central cavity to bind an extended pocket at the B55-PP2Aa interface, positioning mL309_C_ to bind a hydrophobic pocket in PP2Aa (**Extended Data Fig. 11**a-d; residues corresponding to the different subunits of the complex are denoted by subscripts: PP2Aa, A; B55, B; PP2Ac, C; ARPP19, R; FAM122A, F). Overlaying the PP2A:B55-inhibitor complexes with PP2A:B56 (pdbid 2IAE^31^; superimposed using PP2Ac) shows that the PP2Ac mL309_C_ residues are more than 36 Å apart (**Fig. 2e**). This interaction buries ∼1900 Å^2^ of solvent accessible surface area (SASA), explaining the importance of this posttranslational modification for PP2A:B55 complex formation and stability.^19^

The remaining unaccounted density corresponds to *tp*ARPP19 or FAM122A (*tp*ARPP19 residues 42-75 and 86-112; FAM122A residues 81-111). *tp*ARPP19 interacts exclusively with PP2A:B55 subunits B55 and PP2Ac, using helices connected by extended yet ordered loops (helices are pre-populated in free ARPP19, **Extended Data Figs. 4a,c,i)**. This allows *tp*ARPP19 to span the entire front surface of B55 and the PP2Ac active site. Furthermore, in a highly unusual conformation, *tp*ARPP19 loops back on itself to form a stable, overlaid cross at the B55-PP2Ac interface. Together, the interaction buries more than 5200 Å^2^ of solvent accessible surface area (SASA). FAM122A also binds exclusively to B55 and PP2Ac and does so, again, using helices (pre-populated in free FAM122A; **Extended Data Figs. 4d,f**). FAM122A binds a short surface on B55 (the B55 binding platform) and across the PP2Ac active site; the interaction buries 2700 Å^2^ SASA. Notably, both inhibitors bind the most conserved surfaces of B55 (**Fig. 2f**). Despite their common function, these structures show that FAM122A and *tp*ARPP19 bind and inhibit PP2A:B55 using highly distinct mechanisms.

### B55-specific recruitment of tpARPP19

*tp*ARPP19 binds PP2A:B55 using a tripartite mechanism: (a) the N-terminal residues 25-61 bind the top of B55 (B55 site 1), (b) *tp*S62 and the remainder of helix α4 bind PP2Ac and (c) residues 86-112 interact with B55 in a pocket (B55 site 3) nearly 50 Å away from site 1 (**Fig. 3a**). ARPP19 residues 25-41 include helix α2 (25-34) and bind the cleft between B55 loops L4/5 and L5/6 (B55 adopts a WD40 fold composed of 7 β-propellers connected by 7 loops; L1/2 refers to the loop connecting β-propellers 1 and 2). The density for these residues is present but amorphous, suggesting that α2 remains somewhat mobile in the B55 bound state (fuzzy interaction^25,32,33^). Consistently the NMR data show that these residues interact with B55, as the intensities of these N/H^N^ cross-peaks are reduced when bound to either PP2A:B55 or B55_LL._ To test α2 binding to PP2A:B55 in cells, we generated YFP-ARPP19 variants in which either 5 contiguous (5-ala, aa ^32^AAAAA^36, 37^AAAAA^41^) or single amino acids (aa 32-41) were changed to alanine (or A34G) and then tested their ability to pull-down PP2A:B55 from cells (**Fig. 3b; Extended Data Figs. 12a,b**). Although only a single point variant exhibited reduced B55 and PP2Ac binding (Y36A), both 5-ala variants of YFP-ARPP19, 32-36 and 37-41 were unable to pull-down B55. Together, these data show that ARPP19 residues 25-41 contribute to B55 recruitment.

**Figure 3.**
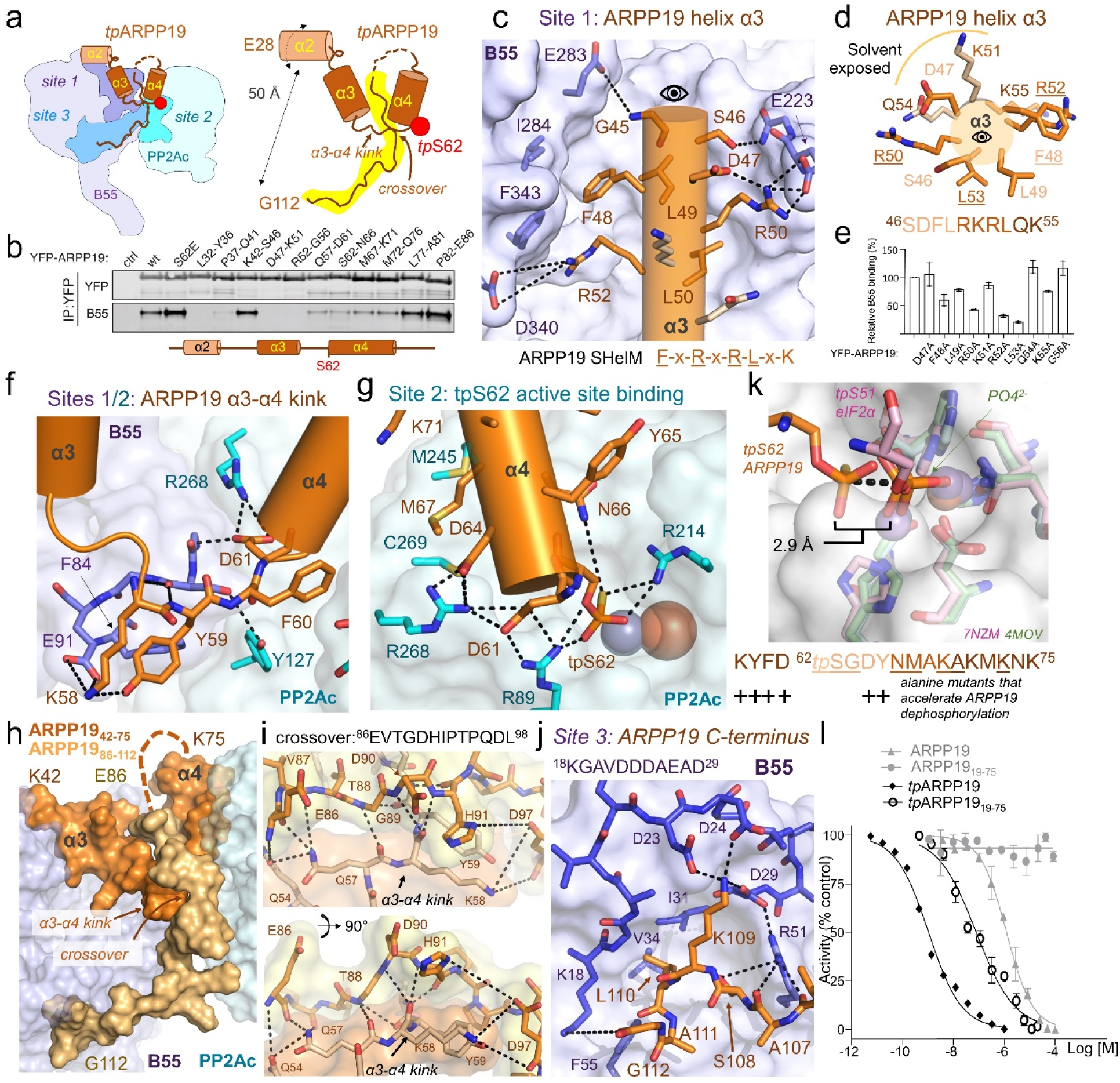
*tp*ARPP19 binding to PP2A:B55. **a**. Cartoon describing the binding of *tp*ARPP19 to PP2A:B55; terminology for the multiple binding sites. **b**. 5 alanine mutational scan through amino acids L32-E86 of ARPP19. The indicated YFP-ARPP19 constructs were transfected into HeLa cells and subsequently immunopurified. Binding efficiency of the YFP-ARPP19 derivatives to B55a was determined by Western blotting. **c**. Site 1: ARPP19 helix α3 interaction with the B55 binding platform. **d**. Helical wheel of ARPP19 helix α3. Residues that interact with B55 are colored orange; solvent exposed residues are light orange. Colors of the labels indicate residue within the same helical turn. Primary sequence colored as helix and key interacting residues are underlined. **e**. Single amino acid change to alanine pull-down analysis of ARPP19 helix α3. **f**. ARPP19 α3-α4 kink; dramatic splaying of Y59_R_, F60_R_ and D61_R_ allow for this conformation to occur. **g**. Phosphorylated S62 binding to the PP2Ac active site. **h**. ARPP19 folds back on itself and binds B55 at site 3. ARPP19 helices α3/α4 are shown in orange; ARPP19 backwards folding is shown in yellow (all surface), highlighting the crossover above the α3-α4 kink (see e.) and the B55 binding site 3 (over 50 Å distant from helix α2). **i**. Intramolecular interactions at the ARPP19 crossover. **j**. ARPP19 binding at B55 binding site 3 **k**. Overlay of the PPP active site residues of PP2A:B55-tpARPP19 (orange, *tp*S62 ARPP19, cyan PP2Ac), PP1:phosphate (4MOV; PP1, green) and a pre-dephosphorylation complex (7NZM; tpS51 eIF2α, pink; PP1, light pink) binds 2.9 Å away from the phosphate moieties of the pre- and post-dephosphorylation complexes; *below,* residues adjacent to S62_R_ that, when mutated, result in faster dephosphorylation of S62^15^. **l**. IC_50_ curves for PP2A:B55 inhibition by different ARPP19 (with and without phosphorylation) constructs. IC_50_ values are reported in **Table 1**. Data are presented as mean values +/- SD, n = 3 experimental replicates.

Following helix α2, the density for *tp*ARPP19 is well-defined, extending down towards the B55 platform, defined by β-propellers 2-4 and loops L1/2, L2/3 and L3/4. These interactions position ARPP19 helix α3 (^46^SDFLRKRLQK^55^) to bind the B55 platform with all ARPP19 residues except D47_R_ and K51_R_ making multiple interactions with B55 (**Fig. 3c**). Because these residues form a helix, amino acids distal in sequence are adjacent in space (**Fig. 3d**). F48_R_ and R52_R_ π-stack and form hydrophobic contacts with B55 loops L4/5 and L5/6 (I284_B_/Y337_B_/F343_B_) while L49_R_ and L53_R_ form hydrophobic contacts with L2/3 and L3/4 (Y178_B_/L198_B_/L225_B_/V228_B_). R50_R_ and R52_R_ also stabilize L3/4 or L5/6, respectively, via a salt bridge (E223_B_/D340_B_) (**Fig. 3c**). The key interactions between the B55 platform and ARPP19 are mediated by ARPP19 residues F-L-R-x-R-L-x-K. Because these residues in ARPP19 are helical, and not extended, we refer to this sequence as a <u>s</u>hort <u>heli</u>cal <u>m</u>otif, or SHelM. To confirm if these ARPP19 residues are important for B55 binding in cells, we generated three 5-ala and 10 single point variants for the ARPP19 α2-α3 loop and helix α3 and tested their ability to pull-down PP2A:B55 form cells. B55 binding to the ^42^AAAAA^46^ variant was unchanged. in contrast, the ^47^AAAAA^51^ and ^52^AAAAA^56^ variants failed to pull-down B55 (**Fig. 3b**). Multiple single alanine mutations for ARPP19 residues 47-56 showed reductions in B55 and PP2Ac binding, particularly L53A and R52A (≤25% vs wt), R50A and F48A (∼50%) and L49A and K55A (∼75%), while no change in B55 binding was observed for D47A, K51A, Q54A and G56A (**Fig, 3e; Extended Data Fig. 12**c). These data are fully consistent with the structure, as mutating residues that interact directly with B55 (F48, L49, R50, R52, L53, K55) reduce binding while those that are mostly solvent accessible (D47, K51, Q54 and G56) do not.

Next, ARPP19 extends towards the B55 L1/2 loop, where it kinks 180° to bind to PP2Ac (**Fig. 3a,f**). At this junction Y59_R_, F60_R_ and D61_R_ are splayed apart, with Y59_R_ binding B55, F60_R_ binding PP2Ac and D61_R_ binding both B55 and PP2Ac. These extended interactions rigidify the ARPP19 backbone, positioning *tp*S62 directly above the PP2Ac active site metals where it forms bipartite salt bridges with the substrate-coordinating arginine residues, R89_C_ and R214_C_ (**Fig. 3g**). This interaction is further stabilized by D61_R_ and D64_R_ which form salt bridges with R268_C_ and R89_C_, resulting in an extended network of ionic interactions between ARPP19 and PP2Ac that stabilize the *tp*S62 conformation. ARPP19 helix α4 (^62^*tp*SGDYNMAKAKMKNK^75^) extends from the PP2Ac active site towards the PP2Ac C-terminus. In this way, ARPP19 helices α3 and α4 interact with B55 (α3) and PP2Ac (α4) using a helix-turn-helix ‘V’ conformation. ARPP19 residues ^76^QLPTAAPD^83^ remain mobile when bound to PP2A:B55. Consistent with the structure, pull-downs using YFP-ARPP19 5-ala variants, in which the kink and helix α4 residues are mutated to alanine (^57^AAAAA^61, 62^AAAAA^66^, ^67^AAAAA^71, 72^AAAAA^76^), weaken B55 binding, albeit not to the same extent as 5-ala variants of α2 or α3 (**Fig. 3b**).

### The ARPP19 crossover

In a highly unusual conformation, the ARPP19 residues following the mobile 76-85 loop back towards the PP2Ac active site. This positions *tp*ARPP19 residues ^86^EVTGDHIPTPQDL^98^ to cross over and stabilize the splayed Y59_R_ and F60_R_ residues at the B55-PP2Ac interface (referred to hereafter as the ‘crossover’, **Figs. 3a,h,i**). This interaction leverages hydrophobic and polar contacts, with E86_R_/V87_R_/T88_R_ binding *tp*ARPP19 (F60_R_/Y65_R_/A66_R_) and I92_R_/P93_R_/P95_R_/L98_R_ binding *tp*ARPP19, PP2Ac and B55 (Y59_R_/F60_R_, V126_C_/Y127_C_/G215_C_ and F84_B_/S89_B_). It is also stabilized by G89_R_, which, because it lacks a Cβ atom, facilitates the close approach needed for backbone hydrogen bonding between the ARPP19 residues ^57^QKYF^60^ and ^89^GDH^91^. Together, these intra-(tpARPP19) and inter-(tpATPP19, PP2Ac, B55) tripartite interactions further stabilize the inhibitory conformation of ARPP19 (**Fig. 3i**). After the crossover, ARPP19 residues ^100^QRKP<u>A</u>L^105^ (S104A prevents phosphorylation of S104 by MASTL; **Extended Data Fig. 4**g) make limited contact with B55. Despite this, they are ordered because the remaining residues, ^106^VASKLAG^112^, especially L110_R_, bind a deep hydrophobic pocket between B55 β-propellers 1 and 2, allowing K109_R_ to interact with multiple acidic residues in B55 loop L7/1 (^22^DDDVAEAD^29^) (**Fig. 3j**). The presence of this ARPP19 binding site on B55, independent of phosphorylation state, is fully consistent with our NMR data (**Figs. 1d,g; Extended Data Figs. 5a-d**).

### ARPP19-mediated inhibition of PP2A:B55

MASTL-phosphorylated ARPP19 is both an inhibitor and substrate of PP2A:B55.^4,6,7,15,18^ To understand why MASTL-phosphorylated ARPP19 is only slowly dephosphorylated by PP2A:B55, we overlayed the structures of PP2A:B55-ARPP19 via its PP2Ac subunit with the PPP subunit of both a PPP product complex (PP1 with a phosphate bound at the active site, PDBID 4MOV^27^) and a PPP pre-dephosphorylation complex (phosphorylated ^p^eIF2α trapped by the catalytically deficient PP1 D64A variant, PDBID 7NZM^34^). The thiophosphate and phosphate in the pre-dephosphorylation and product complexes overlap nearly perfectly. In contrast, the thiophosphoryl group of *tp*ARPP19 is ∼3 Å further away from both metals, in a position that is unproductive for dephosphorylation (**Fig. 3k**). This inhibitory conformation is stabilized by the hydrophobic, polar and ionic interactions between the ARPP19 MASTL recognition residues (^58^KYFDSGDY^65^) and B55 and PP2Ac and the ARPP19 crossover (^86^EVTGDHIPTPQDL^98^), which is secured in place by the interaction of the ARPP19 C-terminus at B55 site 3. Consistent with this, previous data showed that mutating either the MASTL recognition residues (K58A, Y59A, F60A, D61A, D64A, Y65A) or deleting the C-terminus converts ARPP19 into a substrate (**Fig. 3k**; faster dephosphorylation).^15^ Similarly, comparing the IC_50_ values of ARPP19 and a C-terminal deletion (ARPP_19-75_) with and without thiophosphorylation shows that the interaction of the C-terminus with B55 is essential for the potent inhibition of PP2A:B55 (**Fig. 3l**; **Table 1**), as the C-terminal deletion variants either fail to inhibit (non-phosphorylated ARPP19 vs ARPP19_19-75_) or become a >50-fold weaker inhibitor (*tp*ARPP19 vs *tp*ARPP19_19-75_). These overlapped structures also suggest that the mechanism by which ARPP19 ultimately becomes a substrate is that pS62 changes conformation, allowing the phosphate to shift in the active site to a position in which the metal-activated nucleophilic water can mediate dephosphorylation. Our data suggest this is most likely achieved by the release of the ARPP19 C-terminal tail from its interaction with B55.

### Inhibition of PP2A:B55 by FAM122A

The interaction of FAM122A with PP2A:B55 is strikingly different than that of ARPP19, with FAM122A residues 81-111 binding PP2A:B55 using two helices (**Fig. 4a**). Helix α1 (the B55-binding helix) binds B55 while helix α2 (the inhibition helix) binds PP2Ac and blocks the active site. Although FAM122A residues 29-66 were not sufficiently ordered to be modeled, our NMR and binding data suggest that they contribute to binding via a dynamic (fuzzy) charge:charge interaction^25,32,33^ (**Table 2, Extended Data Fig. 3**). The FAM122A B55-binding helix binds the B55 platform, with its N-terminus pointing towards the center of B55 and its C-terminus pointing towards PP2Ac (**Fig. 4b,c**). B55 helix residues L85_F_ and I88_F_ are adjacent in space, allowing them to bind the same shallow hydrophobic pockets on B55 used also by ARPP19 (L49_R_\L53_R_) (**Fig. 4d**). FAM122A residue R84_F_ forms intramolecular polar and ionic interactions with Q87_F_ and E91_F_ that stabilize the short B55 binding helix (**Fig. 4e**). These interactions also allow R84_F_ to form a bidentate salt bridge with D197_B._^35^ Lys89_F_ forms hydrogen bonds with the carbonyls of L3/4 residues E223_B_ and L225_B_ (**Extended Data Fig. 12**d). Finally, E92_F_ binds a deep, basic pocket below the B55 platform where it coordinates residues from 3 B55 loops: L1/2, L2/3 and L3/4 (**Fig. 4f**). The key interactions between B55 and FAM122A are mediated by FAM122A residues R-L-x-x-I-K-x-E-E, four of which (R84_F_, K89_F_, E91_F_ and E92_F_) are highly conserved (**Extended Data Fig. 1**a). Mutating either pair of the B55 helix basic/hydrophobic residues (^84^RLhqIKqEE^92^ to <u>Aa</u>hqIKqEE, ^84^AA^85^, or to Rlhq<u>AA</u>qEE, ^88^AA^89^) residues reduced FAM122A binding by 1.6- and 2.0-fold, respectively (**Fig. 4g**, **Table 2**). Similarly, pull-down assays using PP2A:B55 incubated with FAM122A WT, ^84^AA^85^ and ^88^AA^89^ showed comparable reductions in binding (**Fig. 4h**). Consistent with their weaker affinities, the IC_50_ values of the ^84^AA^85^ and ^88^AA^89^ variants increased by 38- and 48-fold, respectively, (**Table 1**; **Fig. 4g**). Because the FAM122A E92K mutation was identified in cancer tissues (cBioPortal), we also generated the E91K and E92K variants and showed they both bound PP2A:B55 more weakly, with the K_D_s increasing by ∼2-fold. They also inhibited PP2A:B55 more poorly (IC_50_ vs WT: E91K, 18-fold reduction, E92K, 9-fold reduction) (**Tables 1, 2; Extended Data Figs. 2,3**). These data show that E91_F_ and E92_F_ are important for PP2A:B55 binding and inhibition.

**Figure 4.**
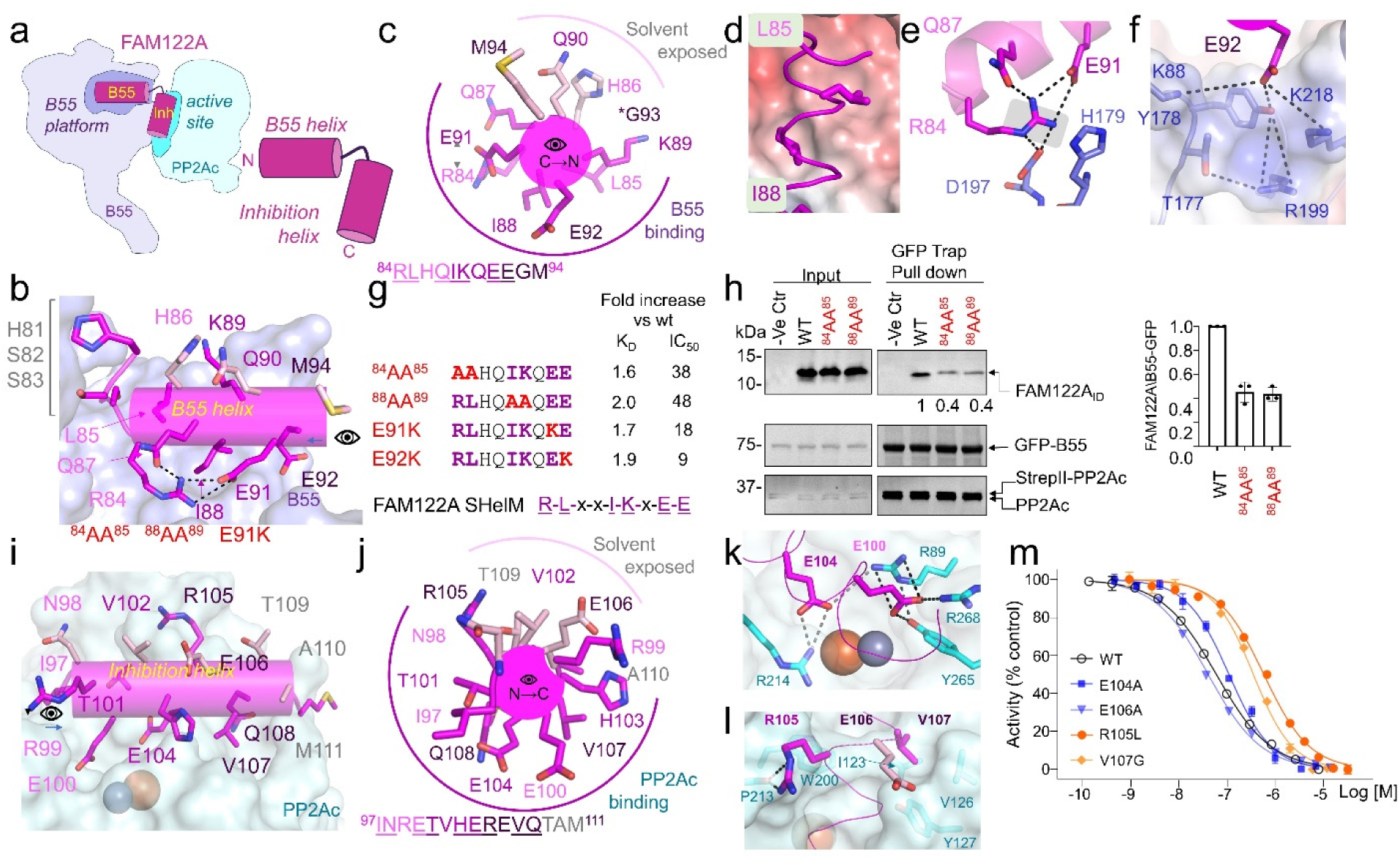
FAM122A binding to PP2A:B55. **a**. Cartoon describing the binding of FAM122A to PP2A:B55. **b**. FAM122A residues R84_F_-E92_F_ form a 3-turn helix when bound to B55. **c**. Helical wheel of the B55 binding helix. Residues that interact with B55 are colored magenta; solvent exposed residues are light pink; colors of the labels indicate residue within the same helical turn. **d.** L85_F_ and I88_F_ bind into shallow pockets on B55 (electrostatic potential surface). **e**. Intra- and intermolecular interactions with Arg84_F_ (polar/ionic interactions shown as dashes). **f**. E92_F_ makes polar/ionic interactions (dashes) with multiple residues from B55. **g**. Mutations in the FAM122A SHelM negatively impact PP2A:B55 binding and inhibition (**Table 2**). **h**. Pulldown assay with SHelM variants shown in (e). Expi293F lysates co-transfected with GFP-B55 and PP2Ac-Strep were incubated with purified PP2Aa and FAM122A variants, pulled-down using a GFP-TRAP and immunoblotted for the indicated proteins. **i**. FAM122A residues Ile97_F_-Met111_F_ form the inhibition helix that binds over the PP2Ac active site. **j**. Helical wheel N→C view of the B55 inhibition helix highlighting residues that interact with PP2Ac and those that are solvent exposed. **k**. Orientation of E104_F_ in the PP2A:B55 active site. **l**. FAM122A residues R105 and V107. **m**. IC_50_ curves for PP2A:B55 inhibition by FAM122A_ID_ E104A, E106A (blue) and R105L, V107G (orange). R105L and V107G are clinical cancer variants (**Table 1**). Data are presented as mean values +/- SD, n = 3 experimental replicates.

Like ARPP19, FAM122A also binds PP2Ac (**Figs. 4a,i**). FAM122A residues C-terminal to the B55 helix form a sharp turn (^94^MDL^96^) and the second helix (α2: ^97^INRETVHEREVQTAM^111^, the inhibition helix) binds PP2Ac, blocking the PP2Ac active site (**Fig. 4j**). This interaction is stabilized by hydrophobic and electrostatic interactions. L96_F_ and I97_F_ bind a hydrophobic pocket adjacent to the PP2Ac active site, which positions E100_F_ to bind Arg268_C_ and R89_C_ and E104_F_ to bind R214_C_ and Arg89_C_ (**Fig. 4k**). The E104A variant only modestly weakens FAM122A inhibition (2-fold; as a control, the inhibitory capacity of E106A, a solvent accessible residue, was not impacted; **Table 1**). These data suggest that interactions at the active site are not essential for PP2Ac inhibition, but instead may be determined by interactions that stabilize its position over the active site. Inhibition helix C-terminal residues Arg105_F_ and Val107_F_, which anchor the helix orientation over the active site, hydrogen bonds with the Pro213_C_ carbonyl and binds a hydrophobic pocket on the opposite side of the helix, respectively (**Fig. 4l**). Interestingly. cBioPortal^36^ highlighted that FAM122A R105L, V107G variants are present in different cancers (FAM122A is a tumor suppressor, as cancer patients that express low levels of FAM122A have significantly worse overall survival than those with high levels of expression^8^). FAM122A R105L and V107G variants showed an 11- and 6-fold less inhibition than WT, respectively (**Fig. 4m**; **Table 1**). These data demonstrate a likely mode of action of these FAM122A cancer variants is that their reduced ability to inhibit PP2A:B55 disrupts PP2A:B55 cellular functions.

### B55-specific recruitment of substrates via the B55 platform

PP2A:B55 specifically dephosphorylates hundreds of substrates,^37,38^ including p107/p130, whose binding domains share sequence similarity with the B55 helix of FAM122A (**Fig. 5a**).^35^ To test if their B55 binding sites overlap, we performed an NMR competition assay (**Fig. 5b**). First, we formed a complex between ^15^N-labeled p107 and B55_LL_ and identified all p107 N/H^N^ cross-peaks that lost intensity due to B55_LL_ binding. We then added an excess of unlabeled FAM122A_Nterm_ and monitored for p107 displacement from B55. All p107 residues that had reduced N/H^N^ cross-peaks intensities due to B55 binding regained their intensities in the presence of excess FAM122A, showing that FAM122A displaced p107 from B55 (**Fig. 5b, Extended Data Figs. 13a-c**). These results establish that, in addition to ARPP19 and FAM122A, p107 and likely more substrates use the B55 platform to bind B55, demonstrating that ARPP19 and FAM122A, in addition to inhibiting the active site, also block substrate binding to PP2A:B55 (**Fig. 5c**).

**Figure 5.**
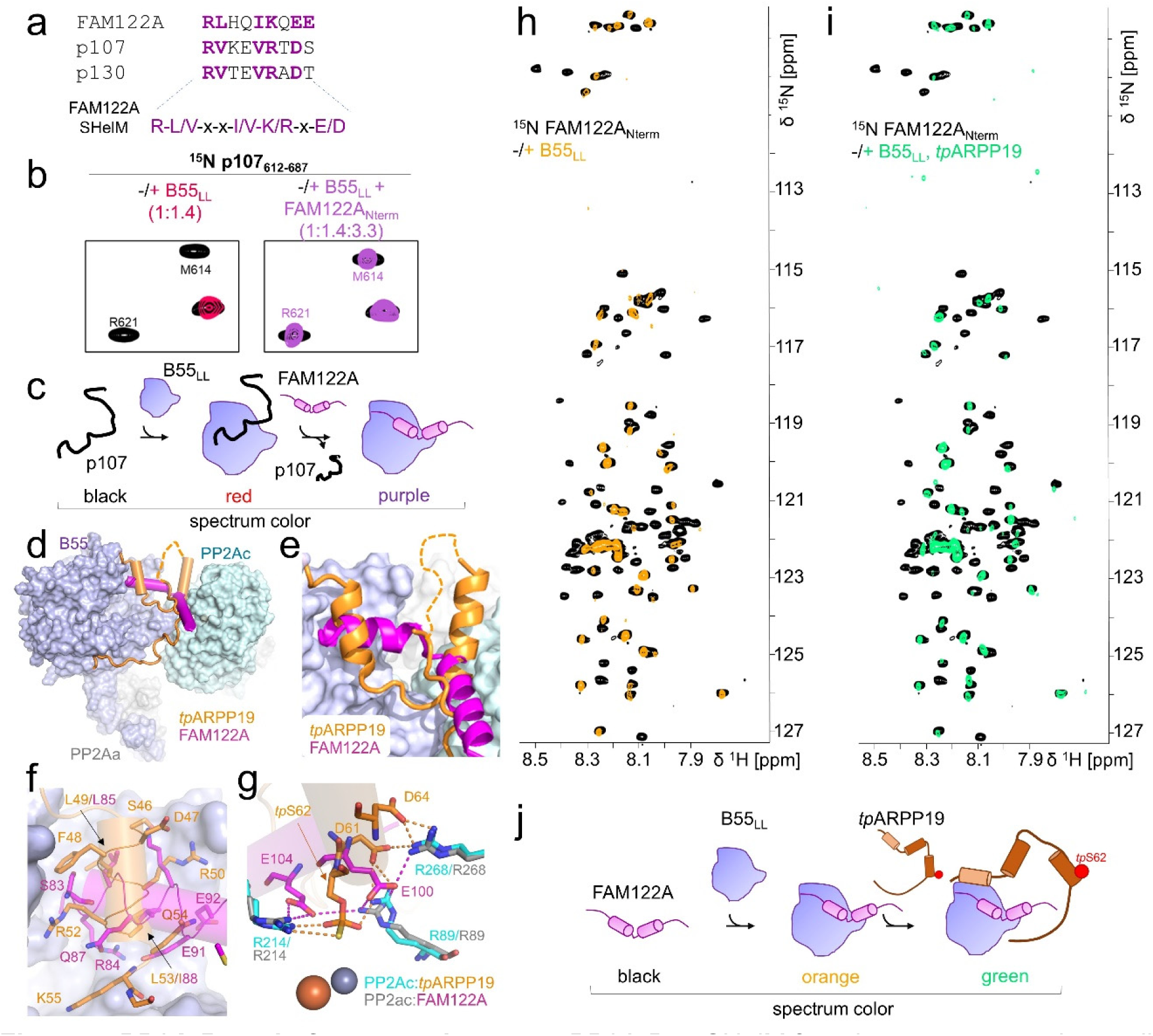
PP2A:B55 platform recruitment. **a**. PP2A-B55 SHelM from interactors experimentally confirmed to bind the B55 platform. **b**. FAM122A displaces p107 from the B55 platform. 2D [^1^H,^15^N] HSQC spectrum of ^15^N-labeled p107 alone (black) and in complex with B55_LL_ (red) shows that specific p107 residues bind B55_LL_. The addition of an excess of unlabeled FAM122A displaces ^15^N-labeled p107 from B55_LL_ and now appears in the 2D [^1^H,^15^N] HSQC spectrum (violet) in an identical position as in the 2D [^1^H,^15^N] HSQC spectrum of ^15^N-labeled p107 alone (black). **c**. Model how FAM122A displaces substrates that bind the B55 platform in a p107 like manner. **d**. Overlay of PP2A:B55 bound to *tp*ARPP19 (orange) or FAM122A (magenta). **e**. Zoom-in of the overlapping regions shown in (d). **f**. Overlay of *tp*ARPP19 (orange) or FAM122A (magenta) on the B55 platform. **g**. Overlay of *tp*ARPP19 (orange) or FAM122A (magenta) at the PP2Ac catalytic pocket (cyan, PP2Ac residues from the ARPP19 complex; grey, PP2Ac residues from the FAM122A complex); ionic interactions indicated by dashed lines. PP2Ac metals are shown as spheres. **h**. 2D [^1^H,^15^N] HSQC spectrum of ^15^N-labeled FAM122A alone (black) and in complex with B55_LL_ (orange). **i**. The addition of unlabeled *tp*ARPP19 does not alter the FAM122A:B55_LL_ complex spectrum (green; identical as orange 2D [^1^H,^15^N] HSQC spectrum in d), highlighting that *tp*ARPP19 and FAM122A can bind simultaneously. **j**. Model of FAM122A and ARPP19 binding B55 simultaneously.

### ARPP19 and FAM122A can bind PP2A:B55 simultaneously

ARPP19 and FAM122A share two PP2A:B55 interaction sites: the B55 platform and the PP2Ac active site (**Figs. 5d, e**). However, their detailed interactions differ. Overlaying both complexes via B55 shows that L53_R_ and I88_F_ bind the B55 platform central hydrophobic pocket while L49_R_ and L85_F_ bind an adjacent, shallower hydrophobic pocket (**Fig. 5f**). The number of intervening residues between these two corresponding hydrophobic amino acids are not identical (4, ARPP19; 3 FAM122A) since the orientations of the bound helices differ by ∼90°, allowing the hydrophobic binding pockets to be engaged distinctly by the two regulators. Similarly, while R52_R_ forms a π-stacking interaction and salt bridge with D340_B_, R85_F_ binds in a pocket nearly 10 Å away, to form a salt bridge with D197_B_. These differences are again due to the distinct binding orientations of these helices. The second shared binding site is the PP2Ac active site (**Fig. 5g**). While both inhibitors use helices to bind, these helices project in opposite directions, with ARPP19 helix α4 extending from the active site towards the PP2Ac C-terminus while the FAM122A inhibitory helix extends to the PP2Ac hydrophobic groove. The only area of overlap is at the active site itself, where *tp*S62_R_ projects deeply into the active site while E100_F_ or E104_F_ bind the edges of the active site pocket.

Outside of these limited shared interactions, ARPP19 and FAM122A bind PP2A:B55 uniquely, with ARPP19 binding B55 at additional interaction sites via helix α2 and its C-terminus. This observation suggests that ARPP19 and FAM122A, which have similar affinities for PP2A:B55 (**Table 2**) may bind PP2A:B55 simultaneously. To test this, we used NMR and pull-down assays. We formed the complex between B55_LL_ and ^15^N-labeled FAM122A_Nterm._ and then added excess of *tp*ARPP19 (**Fig. 5h**). Despite a ∼2.5-fold excess of *tp*ARPP19, no change in the 2D [^1^H,^15^N] HSQC spectrum of the FAM122A_Nterm_ was observed, demonstrating that FAM122A_Nterm_ was not displaced by *tp*ARPP19 (**Fig. 5i**). In an orthogonal experiment, we performed a pull-down competition assay by affinity purifying PP2A:B55 (GFP-B55) in the presence of FAM122A alone or FAM122A with a 5-fold excess of *tp*ARPP19 (**Extended Data Fig. 13**d). Not only were both FAM122A and *tp*ARPP19 pulled down with PP2A:B55, but the amount of FAM122A pulled down in the absence and presence of *tp*ARPP19 was identical. Lastly, we performed the reverse NMR experiment (B55_LL_ bound to ^15^N-labeled ARPP19. and then adding excess unlabeled FAM122A_Nterm_), which showed that ARPP19 stays bound to B55, predominantly via helix α2, in the presence of FAM122A (**Extended Data Fig. 13**e). Together, these experiments show that FAM122A_Nterm_ and *tp*ARPP19 can bind PP2A:B55 simultaneously (**Fig. 5j**) and, thus, that B55 uses its multiple, distinct interaction surfaces to differentially engage B55-specific regulators and/or substrates.

## Discussion

The inhibition of PP2A:B55 by two B55-specific inhibitors, FAM122A and ARPP19, is essential for mitotic entry.^5,8,39,40^ Our data reveal their unexpected, complicated modes of PP2A:B55 binding and inhibition, providing a detailed understanding of their function. Most strikingly, these data show that PP2A:B55 binds its regulators differently than other PPPs. PP1,^25,28,41^ PP2A:B56,^29,42^ calcineurin (PP2B/PP3)^26,43^ and PP4^44^ recruit their cognate regulators and substrates using PPP-specific SLiMs.^45^ In contrast, PP2A:B55 recruits its regulators ARPP19 and FAM122A using α-helices. Different to PPP-SLiM interactions, which are anchored by hydrophobic residues that bind to deep hydrophobic pockets, the B55 platform is comparatively flat with shallow hydrophobic pockets. The lack of pocket depth allows hydrophobic residues to approach and bind via multiple orientations (**Fig. 3**), rather than the single orientation observed in SLiM interactions.^46^ In addition, the B55 platform is bordered by charged residues, especially acidic residues. Because basic residues are long (arginine, lysine), when present in B55 binding helices, they form salt bridges using an array of conformations, as necessitated by the helical binding orientation (**Figs. 3,4**).

Thus, and different from PP2A:B56 or CN where PPP-specific SLiM sequences have been leveraged to identify putative substrates using sequence alone, our data suggests that the analogous strategy is not readily applicable for identifying novel PP2A:B55-specific substrates. Namely, the sequences used by ARPP19 and FAM122A to bind the same pockets of the B55- platform are not highly conserved (ARPP19, F-L-R-x-R-L-x-K; FAM122A, R-L-x-x-I-K-x-E-E) and while some substrates, such as p107/p130, may share similarities with known binding sequences (i.e., FAM122A), our data suggests different B55 substrates may bind the platform via mechanisms not yet observed. Finally, B55 belongs to the WD40 propeller family, and thus adopts a fold that is an established protein interaction domain and demonstrated to bind other proteins using a diverse range of interactions.^47^ Together, these observations, coupled with the discovery that ARPP19 also binds B55 using its C-terminus, suggests that B55 may recruit a set of substrates via interaction surfaces outside the B55 platform used by ARPP19, FAM122A and p107.

In addition to blocking substrate recruitment, our structures also show that FAM122A and *tp*ARPP19 inhibit PP2A:B55 by blocking the PP2Ac active site (**Figs. 3,4**). This combined mechanism of inhibition (blocking substrate recruitment and inhibiting catalytic site access) is also observed in other members of the PPP family, especially PP1. Like FAM122A, Inhibitor-2 (I2) is an IDP inhibitor of PP1^48^ that both blocks PP1-specific substrate/regulator recruitment (by binding PP1-specific SLiM interaction sites, the SILK and the RVxF binding pockets) and blocks catalytic site access (by using a long helix to bind over the PP1 active site in a phosphorylation independent manner; **Extended Data Fig. 14**).^48,49^ This shows that PPP family members PP1 and PP2A:B55 both have endogenous IDP inhibitors that use a common mechanism to potently inhibit their ability to dephosphorylate their cognate substrates, suggesting this may be a mechanism present throughout the PPP family.

Finally, the current literature supports a model in which PP2A:B55 is initially inhibited by FAM122A and later by phosphorylated ARPP19 (**Fig. 1a**),^6–8,18,37^ with the assumption that inhibitor binding is mutually exclusive. However, our NMR and pulldown data show that FAM122A and ARPP19 can bind PP2A:B55 simultaneously, with FAM122A likely binding the B55 platform, and ARPP19, leveraging its multiple B55 interaction sites, binding B55 predominantly via helix α2 in the quaternary complex. The ability of two regulators that share a subset of interaction sites to bind simultaneously to their cognate PPP has also been observed for the PP1:spinophilin:I2 triple complex.^50,51^ In this case, both the PP1:spinophilin and PP1:I-2 heterodimeric complex are stable, with each regulator using the RVxF interaction pocket for PP1 binding.^28,48^ However, they also form a triple complex, in which only spinophilin binds the PP1 RVxF interaction pocket; the I-2 RVxF sequence releases from PP1. It is the extensive interactions of I-2 at the PP1 SILK binding pocket and active site that allows the I-2 RVxF sequence to be dispensable for PP1 binding. Here, we show that ARPP19, FAM122A and PP2A:B55 form a similar complex, in which, in the presence of FAM122A, the interactions of ARPP19 at sites 2 and 3 are dispensable for binding. If and how these ternary interactions contribute to the regulation of PP2A:B55 activity during mitosis remains to be elucidated. This data also suggests that the stable dissociation of FAM122A from PP2A:B55 is needed for formation of the full inhibitory PP2A:B55-*p*ARPP19 complex. One possibility is that a currently unidentified post-translational modification dissociates FAM122A from PP2A:B55 (i.e., phosphorylation; phosphorylation of FAM122A S37 has already been shown to quantitatively dissociate FAM122A from PP2A:B55 to activate the G2/M checkpoint^8^). This would allow for the formation of the full PP2A:B55-*p*ARPP19 inhibitory complex and, once formed, serve as a ‘timer’ to facilitate mitotic exit via pARPP19’s slow transition from an inhibitor to a substrate conformation at the active site. The molecular bases for these events are under active investigation. Together, these studies provide a molecular understanding of regulator and substrate recruitment of the PP2A:B55 holoenzyme. Because of the key regulatory functions of PP2A:B55 in mitosis and DNA damage repair, our data also provides a roadmap for characterizing disease-associated mutations and provide new avenues to therapeutically target this complex, by individually blocking a subset of regulators that use different B55 interaction sites.

## Methods

### Bacterial protein expression

PP2Aa_9-589,_ FAM122A_1-124_ (FAM122A_Nterm_), FAM122A_29-120_ (FAM122A_ID_), FAM122A_67-120_, ARPP19, ARPP19_S104A_, ARPP19_19-75_ and p107_612-687_ were subcloned into pTHMT containing an N-terminal His_6_-tag followed by maltose binding protein (MBP) and a tobacco etch virus (TEV) protease cleavage site. For expression, plasmid DNAs were transformed into *E. coli* BL21 (DE3) RIL or BL21 (DE3) cells (Agilent). Freshly transformed cells were grown at 37°C in LB broth containing kanamycin antibiotics (50 µg/ml) until they reached an optical density (OD_600_) of ∼0.8. Protein expression was induced by addition of 1 mM β-D-thiogalactopyranoside (IPTG) to the culture medium, and cultures were allowed to grow overnight (18-20 hours, 250 rpm shaking) at 18°C. Cells were harvested by centrifugation (8000 *x*g, 15 min, 4°C) and stored at −80°C until purification. Expression of uniformly ^13^C- and/or ^15^N-labeled protein was carried out by growing freshly transformed cells in M9 minimal media containing 4 g/L [^13^C]-D-glucose and/or 1 g/L ^15^NH_4_Cl (Cambridge Isotopes Laboratories) as the sole carbon and nitrogen sources, respectively. FAM122A_ID_ variants E92K, R105L, V107G, S120C, E104A/S120C, E106A/S120C, R84A/L85A/S120C, I88A/K89A/S120C, E91K/S120C, E92K/S120C, FAM_67-120_S120C and ARPP19/S10C were generated by site-directed mutagenesis, sequence verified and expressed as described above.

### Cell culture

Expi293F cells were obtained from ThermoFisher (cat# A14527) and grown in HEK293 Cell Complete Medium (SMM 293-TII, Sino Biological cat# M293TII). For transient overexpression of B55 and PP2Ac constructs, cells were transfected using polyethyleneimine (PEI) transfection reagent. For Western blot and immunoprecipitation studies, whole cell extracts were prepared by lysing cells in ice-cold lysis buffer (20 mM Tris pH 8.0, 500 mM NaCl, 0.5 mM TCEP, 1 mM MnCl_2_, 0.1% Triton-X100, Phosphatase inhibitor cocktail [ThermoFisher]), sonicating and clearing the lysate by centrifuging at 15,000 *x*g for 20 min at 4°C. Total protein concentrations were measured using the Pierce 660 Protein Assay Reagent (ThermoFisher).

### Mammalian protein expression

Full length B55_1-477_ was cloned into pcDNA3.4 including an N-terminal green fluorescence protein (GFP) followed by a TEV cleavage sequence. Full length PP2Ac_1-309_ was cloned into pcDNA3.4 with an N-terminal Strep tag followed by a TEV cleavage sequence. B55 loop-less (B55_LL_), in which B55 residues 126-164 that interact directly with PP2Aa, were removed and replaced with a single NG linker (**Fig. 1B**), was cloned into pcDNA3.4 with an N-terminal green fluorescence protein (GFP) followed by a TEV cleavage sequence. All plasmids were amplified and purified using the NucleoBond Xtra Maxi Plus EF (Macherey-Nagel). B55_WT_ and B55_LL_ were individually expressed in Expi293F cells (ThermoFisher). B55_1-477_ and PP2Ac_1-309_ were co-expressed in Expi293F cells at a 1:2 DNA ratio.

Transfections were performed in 500 mL medium (SMM293-TII, Sino Biological) in 2 L flasks using polyethylenimine (PEI, Polysciences) reagent according to the manufacturer’s protocol in an incubator at 37°C and 8% CO_2_ under shaking (125 rpm). On the day of transfection, the cell density was adjusted to 2.8 X10^6^ cells/mL using fresh SMM293-TII expression medium. DNA of PP2Ac and B55 (2:1 ratio) were diluted in Opti-MEM Reduced Serum Medium (ThermoFisher). Similarly, in a separate tube, PEI 3x the amount of DNA was diluted in the same volume of Opti-MEM Reduced Serum Medium (ThermoFisher). The DNA and PEI mixtures were combined and incubated for 10 min at room temperature, before being added to the cell culture. Valproic acid (2.2 mM final concentration, Sigma) was added to the cells 4 h after transfection and 24 h after transfection sterile-filtered glucose (4.5 mL per 500 mL cell culture, 45%, glucose stock) was added to the cell culture flasks to boost protein production. Cells were harvested 48 hrs after transfection by centrifugation (2,000 x*g* for 20 minutes, 4°C) and stored at −80°C.

### FAM122A purification

Cell pellets expressing FAM122A_Nterm_, FAM122A_ID_ and FAM_67-120_ and variants were resuspended in ice-cold lysis buffer (50 mM Tris pH 8.0, 500 mM NaCl, 5 mM Imidazole, 0.1% Triton X-100, EDTA-free protease inhibitor tablet [ThermoFisher]), lysed by high-pressure cell homogenization (Avestin Emulsiflex C3). Cell debris was pelleted by centrifugation (42,000 x*g*, 45 min, 4°C), and the supernatant was filtered with 0.22 µm syringe filters (Millipore). The proteins were loaded onto a HisTrap HP column (Cytiva) pre-equilibrated with Buffer A (50 mM Tris pH 8.0, 500 mM NaCl, 5 mM imidazole) and eluted using a linear gradient (0-60%) with Buffer B (50 mM Tris pH 8.0, 500 mM NaCl and 500 mM imidazole). Fractions containing the protein were pooled and dialyzed overnight at 4°C with TEV protease (in-house; his_6_-tagged) to cleave the His_6_-MBP tag. Following cleavage, the sample was either (1) loaded under gravity onto Ni^2+^-NTA beads (Prometheus) pre-equilibrated with Buffer A, the flow through and wash A fractions were collected, and then twice heat purified (80°C, 10 min) or (2) twice heat purified (80°C, 10 min). Samples were centrifuged at 15,000 *x*g for 10 min to remove precipitated protein. Supernatant was concentrated and purified using size exclusion chromatography (SEC; Superdex 75 26/60 [Cytiva]) in either NMR buffer (20 mM Na/PO_4_ pH 6.3, 150 mM NaCl, 0.5 mM TCEP), IC_50_ Assay buffer (20 mM Tris pH 8.0, 150 mM NaCl, 0.5 mM TCEP) or fluorescence polarization (FP) Assay buffer (20 mM Tris pH 7.0, 150 mM NaCl, 0.5 mM TCEP). Samples were either directly used for NMR data collection or flash frozen and stored at −80°C.

### ARPP19 Purification

The protocol is identical for all ARPP19 constructs. Cell pellets were resuspended in lysis buffer (50 mM Tris-HCl pH 8.0, 500 mM NaCl, 5 mM Imidazole, 0.1% Triton X-100, EDTA-free protease inhibitor [ThermoFisher]), lysed by high-pressure cell homogenization (Avestin C3-Emulsiflex), cell debris pelleted by centrifugation (42,000 x*g*, 45 min), and the supernatant was filtered with 0.22 µm syringe filters (Millipore). The proteins were loaded onto a HisTrap HP column (Cytiva) pre-equilibrated with Buffer A (50 mM Tris pH 8.0, 500 mM NaCl, 5 mM imidazole), and eluted using a linear gradient (0-60%) of Buffer B (50 mM Tris pH 8.0, 500 mM NaCl, 500 mM imidazole). Fractions containing the protein were pooled and dialyzed overnight at 4°C with TEV protease to cleave the MBP and His_6_-tags. The cleaved protein was incubated with Ni^2+^-NTA resin (Cytiva) and washed with Buffer A. The flow through and Wash A fractions were collected, and heat purified by incubating the samples at 80°C for 20 min. The samples were centrifuged at 15,000 *x*g for 10 min to remove precipitated protein, concentrated and purified using SEC (Superdex 75 26/60 [Cytiva]) in NMR buffer (20 mM Na/PO_4_ pH 6.3, 150 mM NaCl, 0.5 mM TCEP), IC_50_ Assay buffer (20 mM Tris pH 8.0, 150 mM NaCl, 0.5 mM TCEP) or fluorescence polarization (FP) Assay buffer (20 mM HEPES pH 7.0, 150 mM NaCl, 0.25 mM TCEP). Purified samples were again heat purified (80⁰C for 5 min), centrifuged at 15,000 *x*g for 10 min to remove any precipitated protein, and were either directly used for NMR data collection or flash frozen and stored at −80°C.

### PP2Aa purification

Cell pellets expressing PP2Aa_9-589_ were resuspended in ice-cold lysis buffer (50 mM Tris pH 8.0, 500 mM NaCl, 5 mM Imidazole, 0.1% Triton X-100, EDTA-free protease inhibitor tablet [ThermoFisher]), lysed by high-pressure cell homogenization (Avestin Emulsiflex C3). Cell debris was pelleted by centrifugation (42,000 x*g*, 45 min, 4°C), and the supernatant was filtered with 0.22 µm syringe filters. The proteins were loaded onto a HisTrap HP column (Cytiva) pre-equilibrated with Buffer A (50 mM Tris pH 8.0, 500 mM NaCl, 5 mM imidazole) and eluted using a linear gradient (0 to 40%) with Buffer B (50 mM Tris pH 8.0, 500 mM NaCl and 500 mM imidazole). Fractions containing the protein were pooled and dialyzed overnight at 4°C with TEV protease (in-house; his_6_-tagged) to cleave the His_6_-MBP-tag and loaded under gravity onto Ni^2+^- NTA beads (Prometheus) pre-equilibrated with Buffer A. Flow through and wash A fractions were collected, concentrated and loaded onto QTrap HP column (Cytiva) for further purification. The proteins were eluted with a 100 mM – 1 M salt gradient (Buffer A: 20 mM Tris pH 8.0, 100 mM NaCl, 0.5 mM TCEP; Buffer B: 20 mM Tris pH 8.0, 1 M NaCl, 0.5 mM TCEP). PP2Aa fractions were concentrated and further purified using size exclusion chromatography (SEC; Superdex 200 26/60 [Cytiva]) in Assay buffer (20 mM Tris pH 8.0, 150 mM NaCl, 0.5 mM TCEP). Samples were either directly used or flash frozen and stored at −80°C.

### MASTL expression and purification

Expi293F cells were transfected with pcDNA5_FRT_TO_3xFLAG_MASTL as described above. A cell pellet expressing MASTL was resuspended in ice-cold lysis buffer (20 mM Tris pH 8.0, 500 mM NaCl, 0.5 mM TCEP, 0.1% Triton X-100, EDTA-free protease inhibitor tablet [ThermoFisher]), lysed by high-pressure cell homogenization (Avestin Emulsiflex C3). Cell debris was pelleted by centrifugation (42,000 x*g*, 45 min, 4°C), and the supernatant was filtered with 0.22 µm syringe filters (Millipore). Lysates were incubated with Anti-Flag M2 beads (Sigma), pre-equilibrated with Wash Buffer 1 (20 mM Tris pH 8.0, 500 mM NaCl and 0.5 mM TCEP) and slowly rocked at 4°C for 2 hours. Following, beads were washed 3 times with wash buffer (20 mM Tris pH 8.0, 500 mM NaCl, 0.5 mM TCEP, 1 mM MnCl_2_) and bound MASTL protein was eluted by incubating with 150 ng/µL 3x Flag peptide (Biosynthesis) for 10 min. Purified, active MASTL was mixed with 10% glycerol and stored at − 80°C.

### PKA expression and purification

For expression, PKA (human Cα1 in pet15b) was transformed into *E. coli* BL21 (DE3) RIL cells (Agilent). Freshly transformed cells were grown at 37°C in LB broth until they reached an optical density (OD_600_) of ∼0.8. Protein expression was induced by addition of 1 mM β-D-thiogalactopyranoside (IPTG) to the culture medium, and cultures were allowed to grow overnight (18-20 hours, 250 rpm shaking) at 18°C. Cells were harvested by centrifugation (8000 *x*g, 15 min, 4°C) and stored at −80°C until purification. For purification, cell pellets were resuspended in ice-cold lysis buffer (50 mM Tris pH 8.0, 500 mM NaCl, 5 mM Imidazole, 0.1% Triton X-100, EDTA-free protease inhibitor [ThermoFisher]) and lysed by high-pressure cell homogenization (Avestin Emulsiflex C3). Cell debris was pelleted by centrifugation (42,000 x*g*, 45 min, 4°C), and the supernatant was filtered with 0.22 µm syringe filters (Millipore). The proteins were loaded onto a HisTrap HP column (Cytiva) pre-equilibrated with Buffer A (50 mM Tris pH 8.0, 500 mM NaCl, 5 mM imidazole) and eluted using a linear gradient (0-80%) with Buffer B (50 mM Tris pH 8.0, 500 mM NaCl,500 mM imidazole). Fractions containing the protein were pooled and dialyzed overnight in the buffer (20 mM Tris pH 8, 50 mM NaCl, 1 mM EDTA, 2 mM DTT) at 4°C. Purified sample was centrifuged at 15,000 *x*g for 10 min to remove precipitated protein. Supernatant protein sample was mixed with 50% glycerol and stored at −80°C.

### Phosphorylation of ARPP19

Purified ^15^N-labeled-ARPP19 (25 μM) was incubated with either PKA or MASTL kinase (10:1 ratio) in phosphorylation buffer (100 mM Tris pH 7.5, 2 mM DTT, 10 mM MgCl_2_) with 500 µM of ATP-γ-S or ATP (Sigma) for thio-/phosphorylation. The kinase reaction was left at 37⁰C for 72 - 90 hours. Phosphorylated ARPP19 was heat purified by incubating the samples at 80°C for 10 min. The samples were centrifuged at 15,000 *x*g for 10 minutes to remove precipitated kinase and either immediately used for experiments or flash frozen and stored at −80⁰C. Complete phosphorylation was confirmed by chemical shift changes of the phosphorylated serine residue(s) using 2D [^1^H,^15^N] HSQC spectra.

### Immunoprecipitation and Western blot; B55 Vs B55_LL_ interaction with PP2Aa

GFP tagged B55 or B55_LL_ and associated endogenous proteins were captured by incubating equal amounts of total protein (∼500 µg) for each condition with GFP-Trap nanobody agarose beads (prepared using AminoLink Plus Immobilization Kit; ThermoFisher) at 4°C for 16 hours. Following 3 washes with wash buffer (20 mM Tris pH 8.0, 500 mM NaCl, 0.5 mM TCEP, 1 mM MnCl_2_), bound proteins were eluted with 2% SDS sample buffer (90°C, 10 min), resolved by SDS-PAGE (Bio-Rad) and transferred to PVDF membrane for Western blot analysis using indicated antibodies (see reagents table). Purified PP2A:B55 complex was used as a positive control. Antibody fluorescence signals were captured using a ChemiDoc MP Imaging System (Image Lab Touch Software 2.4; Bio-Rad) and band intensities quantified using ImageJ 1.53t.^52,53^

### FAM122A interaction with PP2A:B55 complex

Purified FAM122A and variants (∼25 µg, see preparation in methods) were mixed with Expi293F whole cell extracts expressing B55, PP2Ac constructs and purified PP2Aa. Input samples were collected prior to incubation with agarose beads. GFP-tagged B55 and associated proteins were captured by incubating equal amounts of total protein (∼500 µg) for each condition with GFP-Trap nanobody agarose beads (see preparation in methods) at 4°C for 16 hours. Following 3 washes with wash buffer (20 mM Tris pH 8.0, 500 mM NaCl, 0.5 mM TCEP, 1 mM MnCl_2_), bound proteins were eluted with 2% SDS sample buffer (90°C, 10 min), resolved by SDS-PAGE (Bio-Rad) and transferred to PVDF membrane for Western blot analysis using indicated antibodies (see reagents table). Antibody fluorescence signals were captured using a ChemiDoc MP Imaging System (Image Lab Touch Software 2.4; Bio-Rad) and band intensities quantified using ImageJ 1.53t. Uncropped blots shown in **Extended Data Fig. 15**.

### FAM122A and ARPP19 competition assay

Purified FAM122A_Nterm_ (∼25 µg) and S62 thio-phosphor-ARPP19_S104A_ (∼25 µg or 125 µg, see preparation in methods) alone or in combination were mixed with Expi293F whole cell extracts expressing B55, PP2Ac constructs and purified PP2Aa. Input samples were collected prior to incubation with agarose beads. GFP tagged B55 and associated proteins were captured by incubating equal amounts of total protein (500 µg) for each condition with GFP-Trap nanobody agarose beads (prepared using AminoLink Plus Immobilization Kit; ThermoFisher) at 4°C for 16 hours. Following 3 washes with wash buffer (20 mM Tris pH 8.0, 500 mM NaCl, 0.5mM TCEP, 1mM MnCl_2_), bound proteins were eluted with 2% SDS sample buffer (90°C, 10 min), resolved by SDS-PAGE (Bio-Rad) and transferred to PVDF membrane for Western blot analysis using indicated antibodies (see reagents table). Antibody fluorescence signals were captured using a ChemiDoc MP Imaging System (Image Lab Touch Software 2.4; Bio-Rad) and band intensities quantified using ImageJ 1.53. Uncropped blots shown in **Extended Data Fig. 15**

### PP2Ac methylation – alkaline treatment

For alkaline treatment, 100 μl PP2A:B55 triple complex fraction from anion exchange was mixed with NaOH to a final concentration of 0.2 M and incubated for 10 min at RT. The reaction was neutralized by adding HCl to a final concentration of 0.2 M and diluted to 200 μl with lysis buffer. The control reaction was treated with pre-neutralization solution (0.2 M NaOH and 0.2 M HCl) and diluted to 200 μl with lysis buffer. The samples were boiled with 2% SDS sample buffer (90°C, 10 min), resolved by SDS-PAGE gel (Bio-Rad) and transferred to PVDF membrane for Western blot analysis using indicated antibodies (see reagents table). Antibody fluorescence signals were captured using a ChemiDoc MP Imaging System (Image Lab Touch Software 2.4; Bio-Rad) and band intensities quantified using ImageJ 1.53.

### EGFP_Nanobody protein expression, purification, and immobilization onto agarose beads

For expression, pOPIN_EGFP_Nanobody plasmid DNA (gift from M. Bollen, KU Leuven) was transformed into *E. coli* BL21 (DE3) cells (Agilent). Freshly transformed cells were grown at 37°C in LB broth containing ampicillin antibiotics (50 µg/ml) until they reached an optical density (OD_600_) of ∼0.8. Protein expression was induced by addition of 0.5 mM β-D-thiogalactopyranoside (IPTG) to the culture medium, and cultures were allowed to grow overnight (18-20 hours, 250 rpm shaking) at 18°C. Cells were harvested by centrifugation (8000 *x*g, 15 min, 4°C) and stored at −80°C until purification. Cell pellets expressing EGFP_Nanobody were resuspended in ice-cold lysis buffer (50 mM Tris pH 8.0, 500 mM NaCl, 5 mM Imidazole, 0.1% Triton X-100, EDTA-free protease inhibitor tablet [ThermoFisher]), lysed by high-pressure cell homogenization (Avestin Emulsiflex C3). Cell debris was pelleted by centrifugation (42,000 x*g*, 45 min, 4°C), and the supernatant was filtered with 0.22 µm syringe filters. The proteins were loaded onto a HisTrap HP column (Cytiva) pre-equilibrated with Buffer A (50 mM Tris pH 8.0, 500 mM NaCl, 5 mM imidazole) and eluted using a linear gradient (0-60% B) with buffer B (50 mM Tris pH 8.0, 500 mM NaCl and 500 mM imidazole). Fractions containing the protein were pooled, concentrated, and further purified at room temperature using size exclusion chromatography (SEC; Superdex 75 26/60 [Cytiva]) in PBS pH 7.5 buffer. Purified and concentrated EGFP_Nanobody protein was immobilized onto agarose beads (20 mg protein per column) using AminoLink Plus Immobilization Kit (ThermoFisher), following manufacturer’s instructions in PBS pH 7.5 coupling buffer.

### B55 and B55_LL_ purification

*Expi293F* cell pellets expressing EGFP_B55 or EGFP_B55_LL_ were resuspended in ice-cold lysis buffer (20 mM Tris pH 8.0, 500 mM NaCl, 0.5 mM TCEP, 0.1% Triton X-100, EDTA-free protease inhibitor tablet [ThermoFisher]), lysed by high-pressure cell homogenization (Avestin Emulsiflex C3). Cell debris was pelleted by centrifugation (42,000 x*g*, 45 min, 4°C), and the supernatant was filtered with 0.22 µm syringe filters. Lysates were mixed with GFP-nanobody-coupled agarose beads (see preparation in methods), pre-equilibrated with Wash Buffer 1 (20 mM Tris pH 8.0, 500 mM NaCl and 0.5 mM TCEP) and slowly rocked at 4°C for 2 hours. After two hours, lysate/beads mixture loaded onto gravity columns, the flow through (FT1) was collected and the column was washed 3 times with 25 mL of Wash Buffer (wash 1-3). The GFP-B55 resin was resuspended in 20 mM Tris pH 8.0, 250 mM NaCl and 0.5 mM TCEP, and TEV was added for on-column cleavage rocking overnight at 4°C. The flow through was again collected (FT2) and the resin was washed with 20 mL of Wash buffer 2 (20 mM Tris pH 8.0, 250 mM NaCl and 0.5 mM TCEP; wash 4) and 2x 20 mL with the Wash Buffer 1 (wash 5 and 6). The flow through 2 (FT2) and washes 4-6 were collected, diluted to ∼100 mM salt concentration (with 0 mM NaCl Wash buffer), and loaded onto QTrap HP column (Cytiva) for further purification. The proteins were eluted with a 100 mM – 1 M salt gradient (Buffer A: 20 mM Tris pH 8.0, 100 mM NaCl, 0.5 mM TCEP; Buffer B: 20 mM Tris pH 8.0, 1 M NaCl, 0.5 mM TCEP). B55 or B55_LL_ were concentrated and further purified using SEC (Superdex 200 26/60 [Cytiva]) in NMR buffer (20 mM Na/PO_4_ pH 6.3, 150 mM NaCl, 0.5 mM TCEP) or assay buffer (20 mM Tris pH 8.0, 150 mM NaCl, 0.5 mM TCEP).

### PP2A:B55 complex purification

*Expi293F* cell pellets expressing StrepII_PP2Ac and EGFP_B55 constructs were resuspended in ice-cold lysis buffer (20 mM Tris pH 8.0, 500 mM NaCl, 0.5 mM TCEP, 1 mM MnCl_2_, 0.1% Triton X-100, EDTA-free protease inhibitor tablet [ThermoFisher]), lysed by high-pressure cell homogenization (Avestin Emulsiflex C3). Purified PP2Aa was added to the cell lysate. Cell debris was pelleted by centrifugation (42,000 x*g*, 45 min, 4°C), and the supernatant was filtered with 0.22 µm syringe filters. Lysates were loaded onto a GFP-Nanobody-coupled agarose bead (see preparation in methods) column, pre-equilibrated with Wash Buffer 1 (20 mM Tris pH 8.0, 500 mM NaCl, 1 mM MnCl_2_ and 0.5 mM TCEP) and slowly rocked at 4°C for 2 hours. After two hours, the flow through (FT1) was collected and the column was washed 3 times with 25 mL of Wash Buffer (wash 1-3). The GFP-B55 resin was resuspended in 20 mM Tris pH 8.0, 250 mM NaCl, 1 mM MnCl_2_ and 0.5 mM TCEP, and TEV was added for on-column cleavage rocking overnight at 4°C. The flow through was again collected (FT2) and the resin was washed with 20 mL of Wash Buffer 2 (20 mM Tris pH 8.0, 250 mM NaCl, 1 mM MnCl_2_ and 0.5 mM TCEP) (wash 4) and 2x 20 mL with the Wash Buffer 1 (wash 5 and 6). The flow through 2 (FT2) and washes 4-6 were collected, diluted to ∼100 mM salt concentration (with 0 mM NaCl Wash buffer), and loaded onto Mono Q column (Cytiva) for further purification. The proteins were eluted with a 100 mM – 1 M salt gradient (Buffer A: 20 mM Tris pH 8.0, 100 mM NaCl, 1 mM MnCl_2_ and 0.5 mM TCEP; Buffer B: 20 mM Tris pH 8.0, 1 M NaCl, 1 mM MnCl_2_ and 0.5 mM TCEP). PP2A:B55 complex and B55 fractions were pooled, concentrated and further purified using SEC (Superdex 200 26/60 [Cytiva]) in NMR buffer (20 mM Na/PO_4_ pH 6.3, 150 mM NaCl and 0.5 mM TCEP) or assay buffer (20 mM Tris pH 8.0, 150 mM NaCl, 1 mM MnCl_2_ and 0.5 mM TCEP).

### Cryo-EM data acquisition and processing

The PP2A:B55-FAM122A complex was prepared by purifying PP2A:B55 and incubating it with a 1.5 molar ratio of PP2A:B55-to-FAM122A_ID_ at a total concentration of 1.2 mg/ml. The PP2A:B55-*tp*ARPP19 complex was prepared by purifying PP2A:B55 and incubating it with a 1.5 molar ratio of PP2A:B55-to-*tp*ARPP19 at a total concentration of 2.4 mg/ml. Immediately prior to blotting and vitrification (Vitrobot MK IV, 18°C, 100% relative humidity, blot time 5s), CHAPSO (3-([3-Cholamidopropyl]dimethylammonio)-2-hydroxy-1-propanesulfonate) was added to a final concentration of 0.075% (w/v) for PP2A:B55-FAM122A and 0.125% (w/v) for PP2A:B55-*tp*ARPP19. 3.5 μL of the sample was applied to a freshly glow discharged UltAuFoil 1.2/1.3 300 mesh grid, blotted for 5 s and plunged into liquid ethane. Imaging was performed using a Titan Krios G3i equipped with a Gatan BioQuantum K3 energy filter and camera operating in CDS mode. Acquisition and imaging parameters are given in **Extended Data Table 2**. All data processing steps were performed using Relion 4.0^54^ and are summarized in **Extended Data Figs. 5-8**. For both datasets, micrograph movies were summed and dose-weighted; contrast transfer function (CTF) parameters were estimated using CTFFind 4.1.14^55^ on movie frame-averaged power spectra (∼4 e-/Å^2^ dose). Micrographs were filtered to remove outliers in motion correction and/or CTF estimation results and screened manually to remove micrographs with significant non-vitreous ice contamination. Potential particle locations on the full micrograph set were selected using Topaz^56^ using a model trained on a random subset of the micrographs. Particles on the training subset were selected by a Topaz model trained on previous screening data. Subset picks were subjected to 2D classification, *ab initio* 3D initial model generation, and 3D classification, and surviving particles used to train an improved Topaz model used to pick the full micrograph set. From these picks, 2D classification and 3D classification (with full angular and translational searches) were used to select particles in classes showing clear secondary structure and representing the full complex. Resolution in both datasets was then further improved by cycles of CTF parameter refinement, particle polishing, and fixed-pose 3D classification, alongside the following elaborations: For PP2A:B55-*tp*ARPP19, particles with well resolved ARPP19 density were selected by isolating ARPP19 via signal subtraction of the vast majority of the holoenzyme, followed by fixed-pose 3D classification; this process was performed twice in the course of the processing workflow. The final map was refined from 52,934 particles to a resolution of 2.77 Å. For PP2A:B55-FAM122A, multi-body refinement of the B55 and PP2Ac segments of the complex was needed to resolve details of both segments. Within each resulting body alignment, signal subtraction and fixed-pose 3D classification of FAM122A and its surrounding binding groove was used to select for particles for which multi-body refinement was successful and FAM122A was present and well-resolved. This yielded 103,522 particles for which this was simultaneously true in both bodies. Using these particles, a second multi-body refinement was used to generate maps for model building within each body, with final resolutions of 2.55 Å for the B55 body and 2.69 Å for the PP2Ac body. To generate a consensus map, a refinement was run using only the top 25,000 particles with the smallest sum of squared eigenvalues from the multi-body refinement (as reported by relion_flex_analyse). All 3D auto-refinements for both datasets utilized a soft solvent mask and SIDESPLITTER.^58^ All global map resolutions reported in this work were calculated by the gold-standard half-maps FSC=0.143 metric. Further validation information is given in **Extended Data Figs. 5-8 and Extended Data Table 1**.

### Cryo-EM model building

All models were built and refined by iterating between manual rebuilding and refinement in Coot^58^ and ISOLDE^59^, and automated global real-space refinement in Phenix^60^. For PP2A:B55-FAM122A, the relevant segments of the model were built into the B55 and PP2Ac body maps, using the previously determined crystal PP2A:B55 holoenzyme crystal structure (PDB ID 3DW8) and the available FAM122A AlphaFold model (UniProt Q96E09) as a starting point. The two body models were then joined, and the regions near the joints further rebuilt, and the entire complex refined against the 25,000 particle consensus subset map. For PP2A:B55-*tp*ARPP19, the holoenzyme portion of the PP2A:B55-FAM122A model and the available ARPP19 AlphaFold model (UniProt P56211) were used as starting points. Model geometry and map-model validation metrics are given in **Extended Data Table 1**. Maps in Figure 2 are LAFTER filtered/sharpened maps.^61^

### PP2A:B55 activity assay

Phosphatase activity assays were conducted in 96 well plates (Corning). PP2A:B55 holoenzyme was diluted to desired concentration range (0 to 20 nM) in Enzyme buffer (30 mM HEPES pH 7.0, 150 mM NaCl, 1 mM MnCl_2_, 1 mM DTT, 0.01% triton X-100, 0.1 mg/ml BSA) and incubated at 30°C. The reaction was started by the addition of 6,8-Difluoro 4-Methylumbelliferyl Phosphate (DiFMUP) to a final concentration of 50 μM. Assays were read every 15 s for ∼50 minutes on a CLARIOstarPlus (BMG LABTECH) plate reader (using reader control software version 5.7 R2) and the data was evaluated using GraphPad Prism 9.

### DiFMUP fluorescence intensity assay for PP2A:B55 IC_50_ measurements

DiFMUP based IC_50_ assays were conducted in 384 well plates (Corning, cat# 4411). For ARPP19 and FAM122A IC_50_ assays, PP2A:B55 holoenzyme in Enzyme buffer (30 mM HEPES pH 7.0, 150 mM NaCl, 1 mM MnCl_2_, 1 mM DTT, 0.01% triton X-100, 0.1 mg/ml BSA) was pre-incubated with various concentrations of ARPP19 and FAM122A variants for 30 min at room temperature (**Extended Data Figure 2**). The reaction was started by adding DiFMUP (final concentration 50 μM) into the PP2A:B55-FAM122A enzymatic reaction (final concentration of PP2A:B55 holoenzyme at 1 nM) and then incubated at 30°C for 30 min. End-point reads (ex=360 nm, em=450 nm) were taken on a CLARIOstarPlus (BMG LABTECH) plate reader (using reader control software version 5.7 R2) after the reaction was stopped by the addition of 300 mM potassium phosphate (pH 10). The experiments were independently technically repeated ≥ three times (each reaction was made in n = 3 to 6) and the averaged IC_50_ and standard deviation (SD) values were reported. The data was evaluated using GraphPad Prism 9.

### Fluorescence polarization (FP) PP2A binding assays

Following the instructions of the manufacturer, 100 µM of FAM122A_ID_/S120C (or variants) or ARPP19/S10C was labeled with Alexa Fluor™ 488 C5 Maleimide (ThermoFisher) using 1:10 protein to fluorophore ratio. The mixture was incubated for 2 hours in the dark at room temperature at pH 7.0 and excess β-mercaptoethanol (1.2x the concentration of the fluorophore) was added to inactivate any unreacted Alexa Fluor488. Labeled FAM122A_ID_/S120C (or variants) or ARPP19/S10C was recovered by analytical SEC (Superdex 75 Increase 10/300 [Cytiva]) and used for the fluorescence polarization (FP) assays. The labeled FAM122A_ID_/S120C (or variants) or ARPP19/S10C are henceforth called FAM122A_ID_-tracer, or ARPP19-tracer.

The FP assays were standardized using black 384-well low volume round bottom microplates (Corning, cat# 4411) with 15 µL solution per well. The measurements were performed using a CLARIOstarPlus (BMG LABTECH Inc) microplate reader (using reader control software version 5.7 R2) set up to 482±16 nm excitation, 530±40 nm emission, and dichroic long pass filter 504 nm with reflection ranging between 380-497 nm and transmission ranging between 508-850 nm. For the K_D_ binding measurements, all dilutions were made into fluorescence polarization (FP) buffer (10 mM HEPES pH 7.0, 150 mM NaCl, 0.5 mM TCEP, 0.01% Triton X100, 0.1 mg/mL BSA). A predilution of FAM122A_ID_-tracer/ARPP19-tracer was prepared for 0.3 nM and a serial dilution of PP2A:B55 was made at 3-times the final concentration. 5 µL of FAM122A_ID_-tracer/ARPP19-tracer, 5 µL of serially diluted PP2A:B55 complex and 5 µL of FP buffer were distributed into the 384 well microplate, resulting in a FAM122A_ID_-tracer/ARPP19-tracer 0.1 nM final concentration. All assay experiments were repeated in triplicate and incubated for 30 min in the dark and sealed at room temperature before reading. The experiments were independently repeated ≥ 3-times and the averaged K_D_ and standard deviation values were reported. The data was evaluated using GraphPad Prism 9.

### ARPP19 immunoprecipitation

Synthetic DNA encoding the various ARPP19 sequences was purchased from GeneArt, Life Technologies and cloned into the pcDNA5/FRT/TO (Invitrogen) expression vector containing YFP resulting in YFP-ARPP19 fusion proteins. These constructs were transiently transfected into HeLa cells 24 hours prior to harvesting cells. Cells were lysed in lysis buffer (50 mM Tris-HCl pH 7.5, 50 mM NaCl, 1 mM EDTA, 1 mM DTT and 0.1% NP40). Complexes were immunoprecipitated at 4°C in lysis buffer with GFP-Trap (ChromoTek) beads as described by the manufacturer. Precipitated protein complexes were washed three times in lysis buffer, eluted in 2×SDS sample buffer and subjected to Western blotting using the following antibodies: YFP (1:5000; generated in house), B55α (1:2000; #5689S Cell Signaling Technology), PP2Ac (1:2000; #05-421Millipore). Uncropped blots shown in **Extended Data Fig. 15**

### NMR data collection

All NMR data were collected on either a Bruker Avance Neo 600 MHz or 800 MHz NMR spectrometer equipped with TCI HCN z-gradient cryoprobe at 283 K. (^15^N,^13^C)-labeled FAM122A_Nterm_ (150 µM), (^15^N,^13^C)-labeled FAM122A_ID_ (400 µM), (^15^N,^13^C)-labeled ARPP19 (400 µM) and (^15^N,^13^C)-labeled *pS62pS104*ARPP19 (200 µM) were prepared in either FAM122A or ARPP19 NMR buffer with 5-10% (v/v) D_2_O added immediately prior to data acquisition. The sequence-specific backbone assignments both proteins were determined by recording a suite of heteronuclear NMR spectra: 2D [^1^H,^15^N] HSQC, 3D HNCA, 3D HN(CO)CA, 3D HNCACB, 3D CBCA(CO)NH, 3D HNCO, and 3D HN(CA)CO, with an additional spectrum, 3D (H)CC(CO)NH, collected for FAM122A_ID_ (t_m_ = 12 ms)^62^. Spectra were processed in Topspin (Bruker Topspin 4.1.3) and referenced to internal DSS.

### Sequence-specific backbone assignment, chemical shift index and chemical shift perturbation

Peak picking and sequence-specific backbone assignment were performed using the program CARA 1.9.1 (http://www.cara.nmr.ch). Chemical shift index (CSI) calculations of FAM122A_Ntern_, FAM122A_ID_, ARPP19 and *pS62pS104*ARPP19 were performed using both Cα and Cβ chemical shifts for each assigned amino acid, omitting glycine, against the RefDB database^63^. Secondary structure propensity (SSP) scores were calculated using a weighted average of seven residues to minimize contributions from chemical shifts of residues that are poor measures of secondary structure.^64^ The changes in peak position between different FAM122A or ARPP19 constructs/variants were traced according to nearest neighbor analysis. Chemical shift differences (Δδ) were calculated using the following equation:

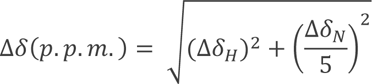

### NMR interaction studies of FAM122A/ARPP19 with PP2A:B55 and B55_LL_

All NMR interaction data of FAM122A_Nterm/ID_ or ARPP19/*pS62pS104*ARPP19 with either PP2A:B55 or B55_LL_ were recorded using a Bruker Neo 600 MHz NMR spectrometer equipped with a HCN TCI active z-gradient cryoprobe at 283 K. All NMR measurements of FAM122A_Nterm_/FAM122A_ID_ or ARPP19 and *pS62pS104*ARPP19 were recorded using ^15^N-labeled protein in NMR buffer and 90% H_2_O/10% D_2_O. For each interaction, an excess of unlabeled B55_LL_ of PP2A:B55 complex (min 25% surplus ratio) was added to the ^15^N-labeled FAM122A/ARPP19 construct under investigation and incubated on ice for 10 minutes before the 2D [^1^H,^15^N] HSQC spectrum was collected. FAM122A/ARPP19 concentrations ranged from 2-6 μM. NMR data were processed using nmrPipe^65^ and the intensity data were analyzed in Poky.^66^ Each data set was normalized to its respective most intense peak and the difference between each free 2D [^1^H,^15^N] HSQC spectrum FAM122A/ARPP19 residue was compared to its respective peak, if present, on the 2D [^1^H,^15^N] HSQC spectrum of FAM122A/ARPP19 in complex with B55_LL_ or PP2A:B55. Any overlapping peaks were omitted for this analysis.

## Data Availability

The NMR data generated in this study have been deposited in the BioMagResBank database under accession code BMRB 51828 (https://bmrb.io/data_library/summary/?bmrbId=51828) (FAM122A_Nterm_), 51682 (https://bmrb.io/data_library/summary/?bmrbId=51682) (FAM122A_ID_), 51881 (https://bmrb.io/data_library/summary/?bmrbId=51881) (ARPP19) and 51882 (https://bmrb.io/data_library/summary/?bmrbId=51882) (*tpS62tpS104*ARPP19). The atomic coordinates and structure factors for PP2A:B55-*tp*ARPP19 complex have been deposited in the PDB database under accession code 8TTB (10.2210/pdb8ttb/pdb) and EMDB code EMD-41604. The atomic coordinates and structure factors for PP2A:B55-FAM122A complex have been deposited in the PDB database under accession code 8SO0 (10.2210/pdb8so0/pdb) and EMDB code EMD-40644. All IC_50_, FP and pull-down data generated in this study are provided in the Supplementary Information and/or Source Data file, which is available at Figshare (10.6084/m9.figshare.23992656).

## Acknowledgements

We thank Drs. Xavier Grana (Temple University), Arminja Kettenbach (Dartmouth College) and Roland Dunbrack (Fox Chase Cancer Center) for fruitful discussions. We thank Dr. André Da Silva Santiago for his help with the project. This work was supported by grant 1R01GM134683 from the National Institute of General Medicine and 1R01NS124666 from the National Institute of Neurological Disorders and Stroke to WP and grant 1R01GM144379 from the National Institute of General Medicine to RP. The work at the Novo Nordisk Foundation Center for Protein Research is supported by grant NNF14CC0001. JBH was funded by grant NNF17OC0025404 from the Novo Nordisk Foundation and the Stanford Bio-X Program. A portion of this research was supported by NIH grant U24GM129547 and performed at the PNCC at OHSU (Dr. Vamseedhar Rayaprolu) and accessed through EMSL (grid.436923.9), a DOE Office of Science User Facility sponsored by the Office of Biological and Environmental Research. Cryo-EM data acquisition was also performed at The University of Chicago Advanced Electron Microscopy Core Facility (RRID:SCR_019198).

## Author Contributions

RP, WP and SKRP developed the concept. SKRP, MRV, RJG expressed and purified all proteins. MRV and WP performed and analyzed NMR experiments. SKRP, JRF, RP determined the cryo-EM structures. SKRP performed pulldown and IC_50_ work. SKRP and RJG performed all fluorescence polarization binding experiments. JBH, TK, JN performed ARPP19 YFP pulldowns. RP, WP, SKRP, MRV, RJG, JRF wrote the manuscript with comments and input from all co-authors.

