## Supplemental Data for "Cryo-EM structures of PP2A:B55-FAM122A and PP2A:B55-ARPP19"

**Table of Contents.**

| | |
|---|---|
| **Extended Data Table 1.** | Model statistics |
| **Extended Data Table 2.** | Cryo-EM image acquisition parameters |
| **Extended Data Figure 1.** | Constructs and PP2A:B55-inhibitor complex formation. |
| **Extended Data Figure 2.** | $IC_{50}$ inhibition assay for ARPP19 and FAM122A variants vs PP2A:B55. |
| **Extended Data Figure 3.** | ARPP19 and FAM122A variants vs PP2A:B55/$B55_{LL}$ binding experiments**.** |
| **Extended Data Figure 4.** | HSQC and CSI plots of ARPP19, FAM122A and MASTL-phosphorylated ARPP19**.** |
| **Extended Data Figure 5.** | NMR data supporting the interaction of phosphorylated ARPP19 and FAM122A with PP2A:B55 |
| **Extended Data Figure 6.** | NMR data supporting the interaction of phosphorylated ARPP19 and FAM122A with $B55_{LL}$. |
| **Extended Data Figure 7.** | Cryo-EM data acquisition and image processing of the PP2A:B55-*tp*ARPP19 complex. |
| **Extended Data Figure 8.** | Cryo-EM 2D class averages and maps of the PP2A:B55-*tp*ARPP19 complex |
| **Extended Data Figure 9.** | Cryo-EM data acquisition and image processing of the PP2A:B55-FAM122A complex |
| **Extended Data Figure 10.** | Cryo-EM 2D class averages and maps of the PP2A:B55-FAM122A complex |
| **Extended Data Figure 11.** | PP2Ac C-terminus interaction with PP2Aa and B55. |
| **Extended Data Figure 12.** | Pull-down studies of ARPP19 with PP2A and the interaction of FAM122A E92 with B55. |
| **Extended Data Figure 13.** | NMR displacement and pull-down studies show that FAM122A displaces p107 and that FAM122A and *tp*ARPP19 can bind simultaneously. |
| **Extended Data Figure 14.** | Overlay of PP2A:B55-*tp*ARPP19, PP2A:B55-FAM122A and PP1:I2. |
| **Extended Data Figure 15.** | Uncropped gel and western blot images. |

**Extended Data Table 1: Model statistics for the cryo-EM structure of PP2A:B55-FAM122 and PP2A:B55-*tp*ARPP19**

| | FAM122A | | | ARPP19 |
|---|---|---|---|---|
| | *B55 body* | *Catalytic body* | *Consensus* | |
| **Composition (#)** | | | | |
| Chains | 4 | 4 | 4 | 4 |
| Atoms | 13319 (Hydrogens: 6636) | 8102 (Hydrogens: 4029) | 21418 (Hydrogens: 10662) | 21926 (Hydrogens: 10924) |
| Residues | 833 | 510 | 1343 | 1376 |
| Ligands | None | ZN: 1 FE: 1 | ZN: 1 FE: 1 | ZN: 1 FE: 1 |
| **Bonds (RMSD)** | | | | |
| Length (Å) (# > 4σ) | 0.003 (0) | 0.002 (0) | 0.002 (0) | 0.002 (0) |
| Angles (°) (# > 4 σ) | 0.463 (0) | 0.460 (0) | 0.400 (0) | 0.398 (0) |
| Dihedral angles (°) (# > 4 σ) | 10.110 (0) | 9.781 (0) | 8.076 (0) | 8.016 (1) |
| **MolProbity score** | 0.81 | 0.88 | 0.95 | 1.15 |
| **Clash score** | 1.05 | 1.24 | 1.87 | 3.65 |
| **EMRinger score** | 5.13 | 4.65 | 4.67 | 5.03 |
| **Ramachandran plot (%)** | | | | |
| Outliers | 0.00 | 0.00 | 0.00 | 0.00 |
| Allowed | 0.49 | 2.18 | 1.89 | 1.70 |
| Favored | 99.51 | 97.82 | 98.11 | 98.30 |
| **Rama-Z (Ramachandran plot Z-score, RMSD)** | | | | |
| whole | 1.05 (0.31) | 1.15 (0.38) | 1.96 (0.24) | 1.83 (0.24) |
| helix | 1.17 (0.30) | 1.17 (0.31) | 1.99 (0.22) | 1.97 (0.23) |
| sheet | 0.47 (0.36) | 0.20 (0.90) | 0.87 (0.37) | 0.87 (0.38) |
| loop | 0.50 (0.39) | 0.43 (0.48) | 0.73 (0.29) | 0.61 (0.28) |
| **Rotamer outliers (%)** | 0.13 | 0.67 | 0.08 | 0.00 |
| **Cβ outliers (%)** | 0.00 | 0.00 | 0.00 | 0.00 |
| **Peptide plane (%)** | | | | |
| Cis proline/general | 0.0/0.0 | 10.0/0.0 | 5.7/0.0 | 5.2/0.0 |
| Twisted proline/general | 0.0/0.0 | 0.0/0.0 | 0.0/0.0 | 0.0/0.0 |
| **CaBLAM outliers (%)** | 0.12 | 2.01 | 0.76 | 0.52 |

| | | | | |
|---|---|---|---|---|
| **ADP/B-factors (min/max/mean)** | | | | |
| Protein | 27.60/179.51/89.51 | 33.76/188.61/121.80 | 8.88/155.34/78.91 | 41.46/188.26/86.61 |
| Ligand | N/A | 162.24/187.03/174.63 | 113.84/135.82/124.83 | 98.59/113.27/105.93 |
| **Map resolution (FSC = 0.143) (Å)** | 2.55 | 2.69 | 2.80 | 2.77 |
| **Map-model fit** | | | | |
| FSC = 0.5 (Å) | 2.61 | 3.12 | 2.86 | 2.81 |
| Masked cross correlation | 0.87 | 0.80 | 0.87 | 0.87 |

**Extended Data Table 2: Cryo-EM image acquisition parameters**

| | **PP2A:B55-FAM122A** | **PP2a:B55-*tp*ARPP19** |
|---|---|---|
| Microscope | Titan Krios G3i, BioQuantum K3 camera | |
| Accelerating voltage | 300 kV | |
| Nominal magnification | 81,000x | 105,000x |
| Calibrated pixel size | 1.068 Å | 0.827 Å |
| Camera mode | Counted super-resolution, CDS | Counted super-resolution, hardware bin 2, CDS |
| Total exposure dose | 70 $e^{-}$/ $Å^2$ | |
| Exposure movie frames | 59 | 62 |
| Energy filter width | 20 eV | |
| Target defocus range | -1.9 to -0.9 µm | |

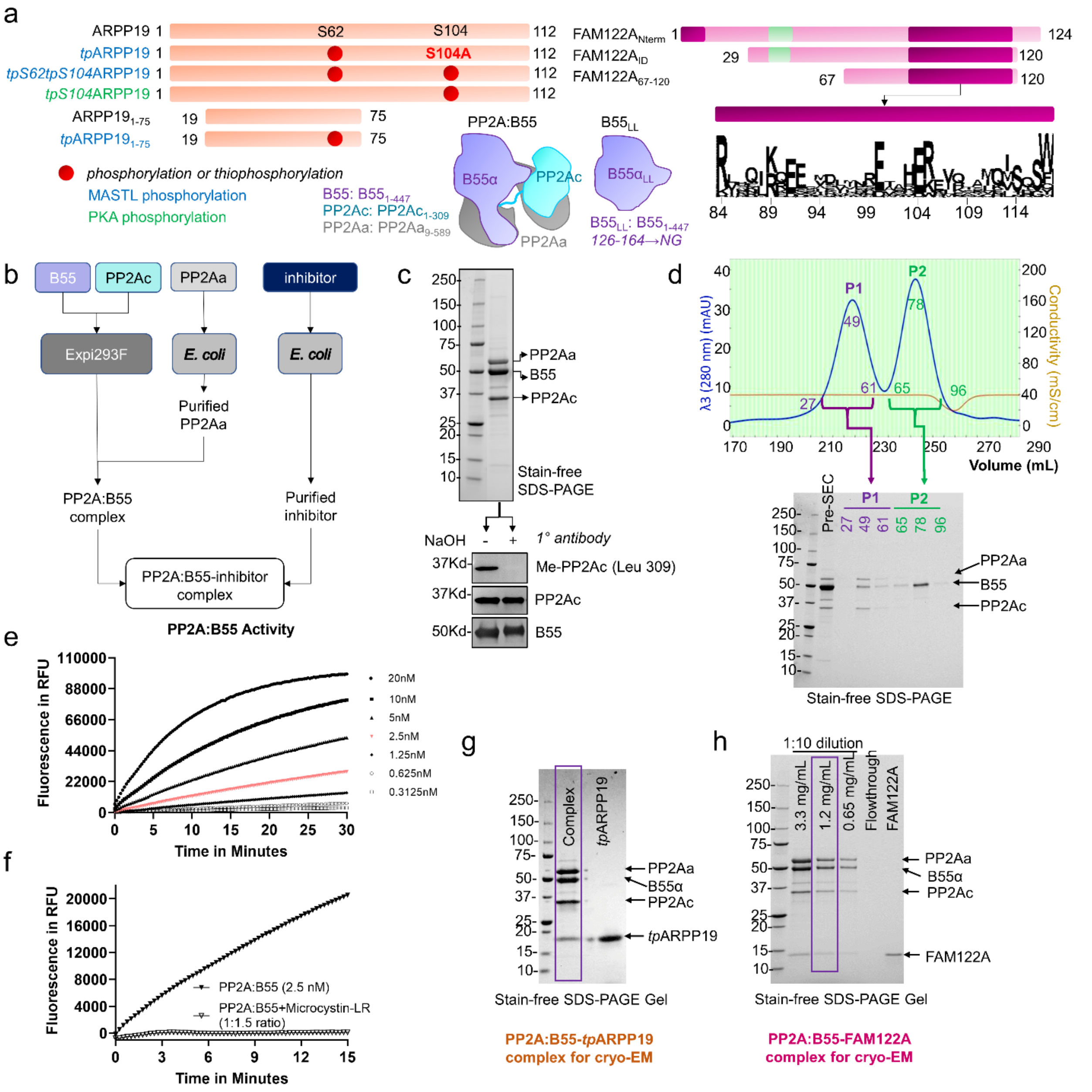

**Extended Data Fig. 1. Constructs and complex production**. **a**. Construct schematic. ARPP19 constructs phosphorylated as indicated. Most highly conserved FAM122A residues are shown. **b**. Schematic describing the production of PP2A:B55 and PP2A:B55-inhibitors for structural and biophysical studies. **c**. Immunoblots of purified PP2A:B55 (above, SDS-PAGE), with and without NaOH treatment, using antibodies detecting methylated PP2Ac (BioLegend, Cat# 828801), PP2Ac (Millipore, Cat# MABE1783) and B55 (Cell Signaling Technologies, Cat# 2290S). **d**. Size exclusion chromatography (SEC) chromatogram of PP2A:B55; peak 1 corresponds to PP2A:B55; peak 2 is excess free B55. **e**. PP2A:B55 activity assay using DiFMUP as a substrate; PP2A:B55 concentrations 0.3125-20 nM; 2.5 nM concentration highlighted in red. **f**. Same as d (2.5 nM concentration) with and without the PPP inhibitor microcystin-LR (3.75 nM concentration). **g**. SDS-PAGE of the PP2A:B55-*tp*ARPP19 used for Cryo-EM grid preparation and data collection. **h**. SDS-PAGE of the PP2A:B55-FAM122A used for Cryo-EM grid preparation and data collection.

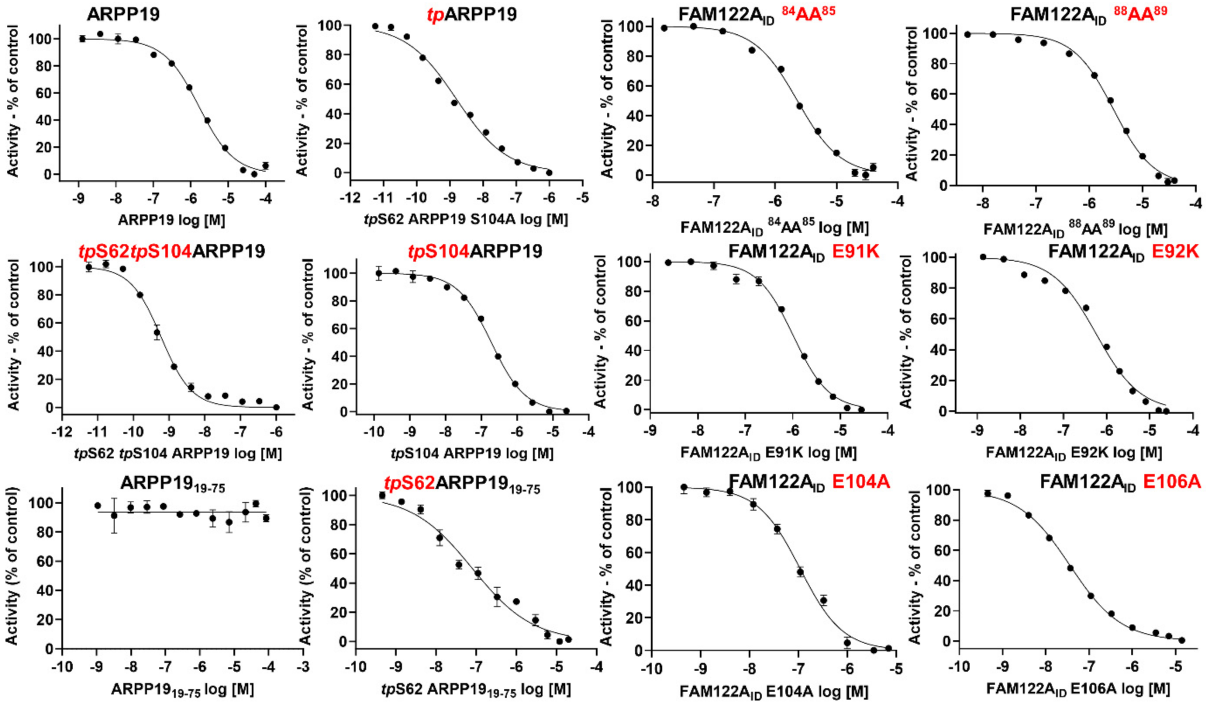

**Extended Data Fig. 2. $IC_{50}$ inhibition assay for ARPP19 and FAM122A variants vs PP2A:B55.** $IC_{50}$ curves for PP2A:B55 inhibition by ARPP19 (variants; unphosphorylated and phosphorylated as well as different constructs) and $FAM122A_{ID}$ variants. $IC_{50}$ values are reported in **Table 1**. Data are presented as mean values ± SD, n = 3 experimental replicates.

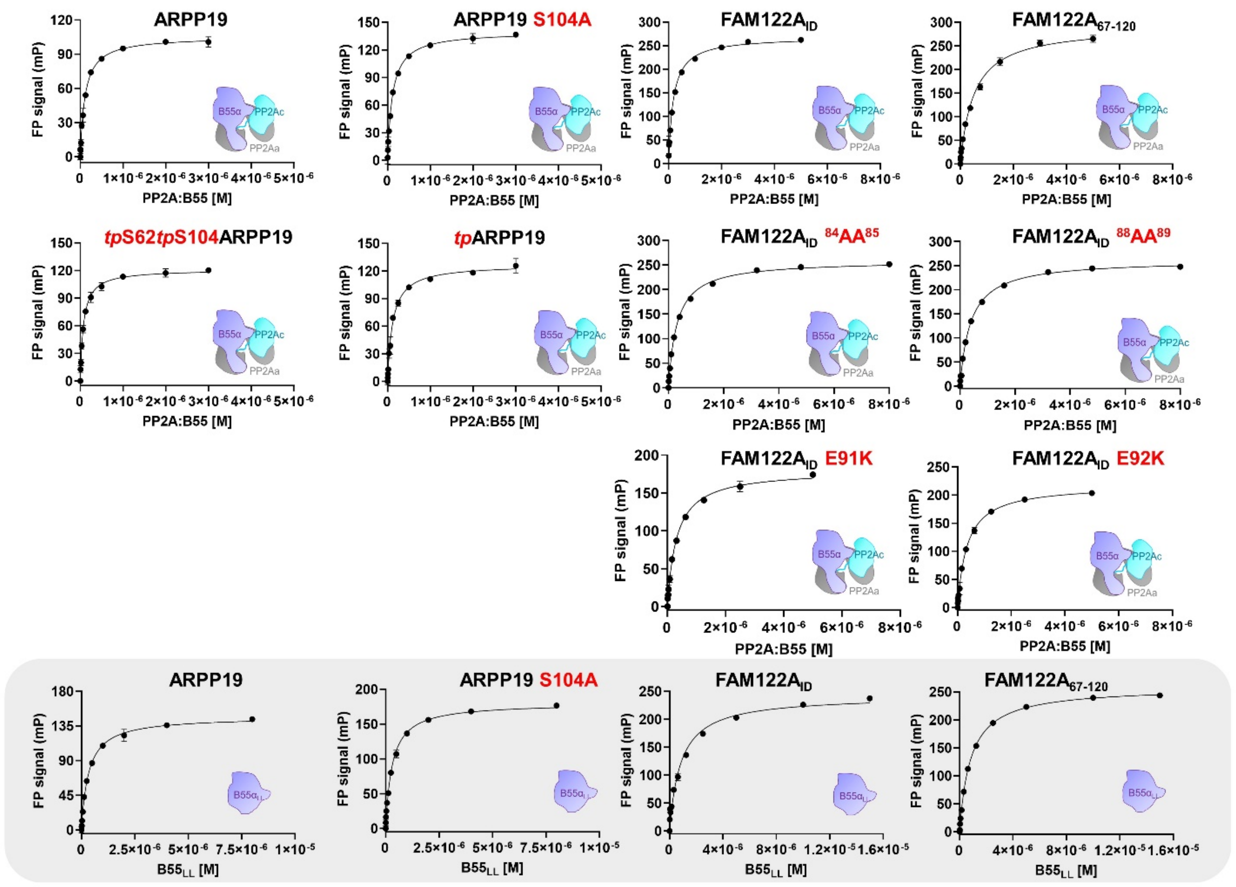

**Extended Data Fig. 3. ARPP19 and FAM122A variants vs PP2A:B55/B55$_{LL}$ binding experiments.** FP binding studies to measure the interaction of ARPP19 (unphosphorylated and phosphorylated) and FAM122A$_{ID}$ variants with PP2A:B55 and B55$_{LL}$ (gray box). K$_D$ values are reported in **Table 2**. Data are presented as mean values ± SD, n = 3 experimental replicates.

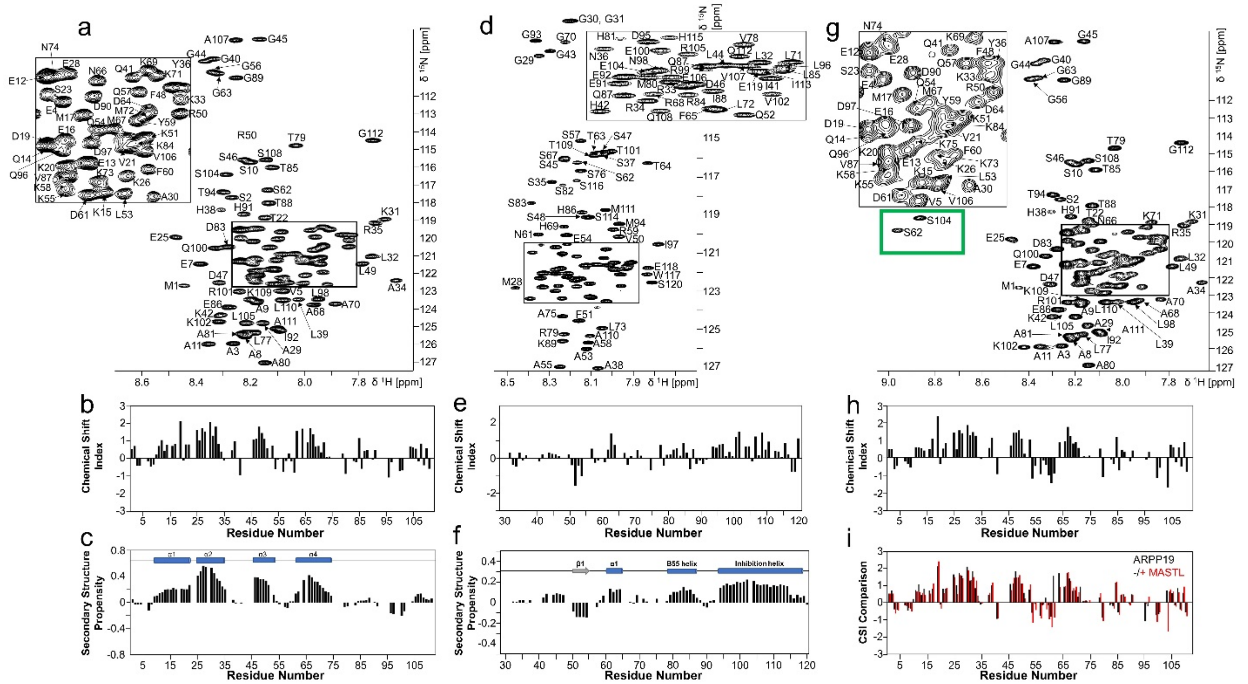

**Extended Data Fig. 4**. **HSQC and CSI plots of ARPP19, FAM122A and MASTL-phosphorylated ARPP19**. **a**. Fully annotated 2D [$^{1}$H,$^{15}$N] HSQC spectrum of $^{15}$N-labeled ARPP19. **b**. Chemical Shift Index (CSI) and **c.** Secondary-structure propensity (SSP) data for ARPP19 plotted vs. residue numbers. (SSP > 0, α helix; SSP < 0, β strand). Cα and Cβ chemical shifts were used to create the CSI and SSP plots (RefDB database). Preferred secondary structure indicated above SSP data. **d**. Fully annotated 2D [$^{1}$H,$^{15}$N] HSQC spectrum of $^{15}$N-labeled FAM122A$_{ID}$. **e**. CSI and **f.** SSP data for FAM122A$_{ID}$ plotted vs. residue numbers; same as c. **g**. Fully annotated 2D [$^{1}$H,$^{15}$N] HSQC spectrum of $^{15}$N-labeled *tpS62tpS104*ARPP19. **h**. CSI; same as c. **i.** CSI comparison between ARPP19 (black) and *tpS62tpS104*ARPP19 (red).

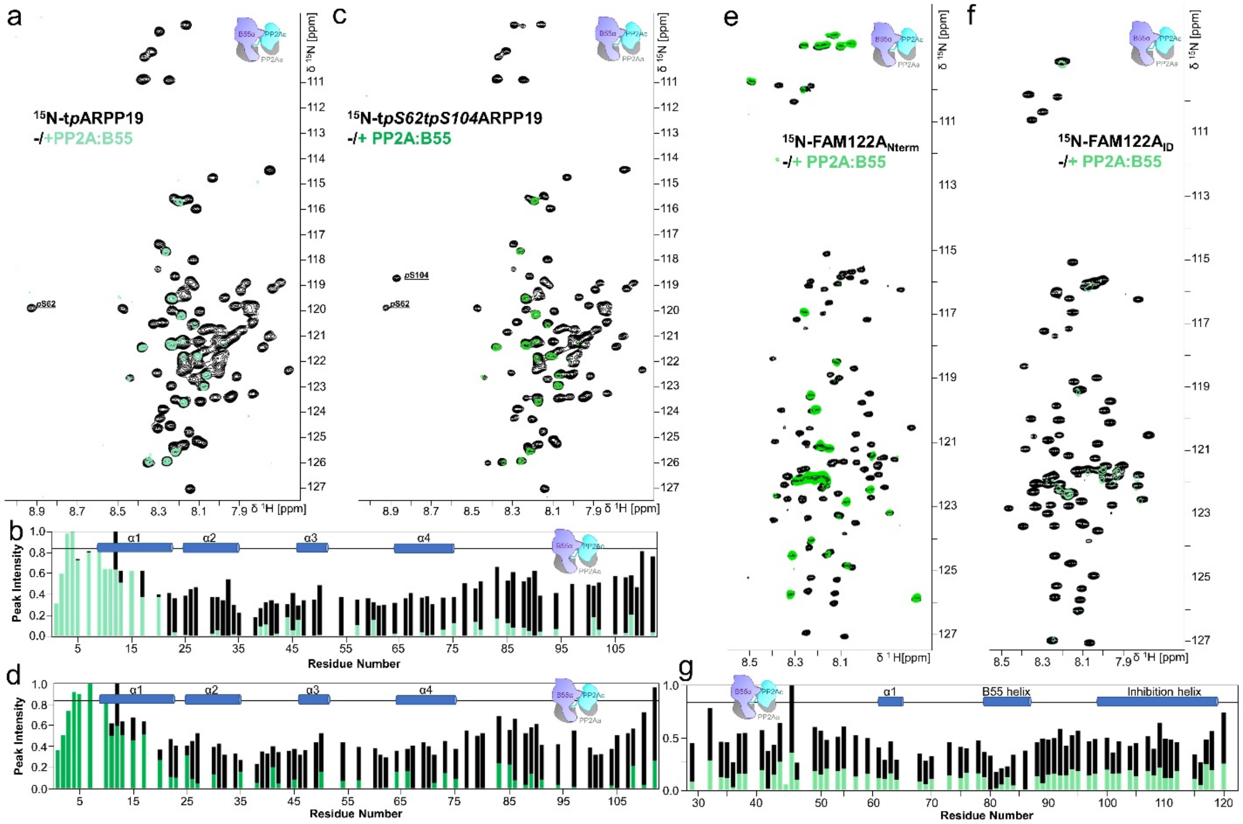

**Extended Data Fig. 5**. **NMR data supporting the interaction of phosphorylated ARPP19 and FAM122A with PP2A:B55**. **a**. 2D [$^1$H,$^{15}$N] HSQC spectrum of $^{15}$N-labeled *tp*ARPP19 alone (black) and in complex with PP2A:B55 (green). pS62 labeled for clarity. **b**. Peak intensity vs ARPP19 protein sequence plot for *tp*ARPP19 alone (black) and when bound to PP2A:B55 (green). Secondary structure elements based on NMR CSI data are indicated. **c**. 2D [$^1$H,$^{15}$N] HSQC spectrum of $^{15}$N-labeled *tpS62tpS104*ARPP19 alone (black) and in complex with PP2A:B55 (green). pS62 and pS104 labeled for clarity. **d**. Peak intensity vs ARPP19 protein sequence plot for *tpS62tpS104*ARPP19 alone (black) and when bound to PP2A:B55 (green). Secondary structure elements based on NMR CSI data are indicated. **e**. 2D [$^1$H,$^{15}$N] HSQC spectrum of $^{15}$N-labeled FAM122A$_{Nterm}$ alone (black) and in complex with PP2A:B55 (green). **f**. 2D [$^1$H,$^{15}$N] HSQC spectrum of $^{15}$N-labeled FAM122A$_{ID}$ alone (black) and in complex with PP2A:B55 (green). **g**. Peak intensity vs FAM122A$_{ID}$ protein sequence plot for FAM122A$_{ID}$ alone (black) and when bound to PP2A:B55 (green). Secondary structure elements based on NMR CSI data are indicated.

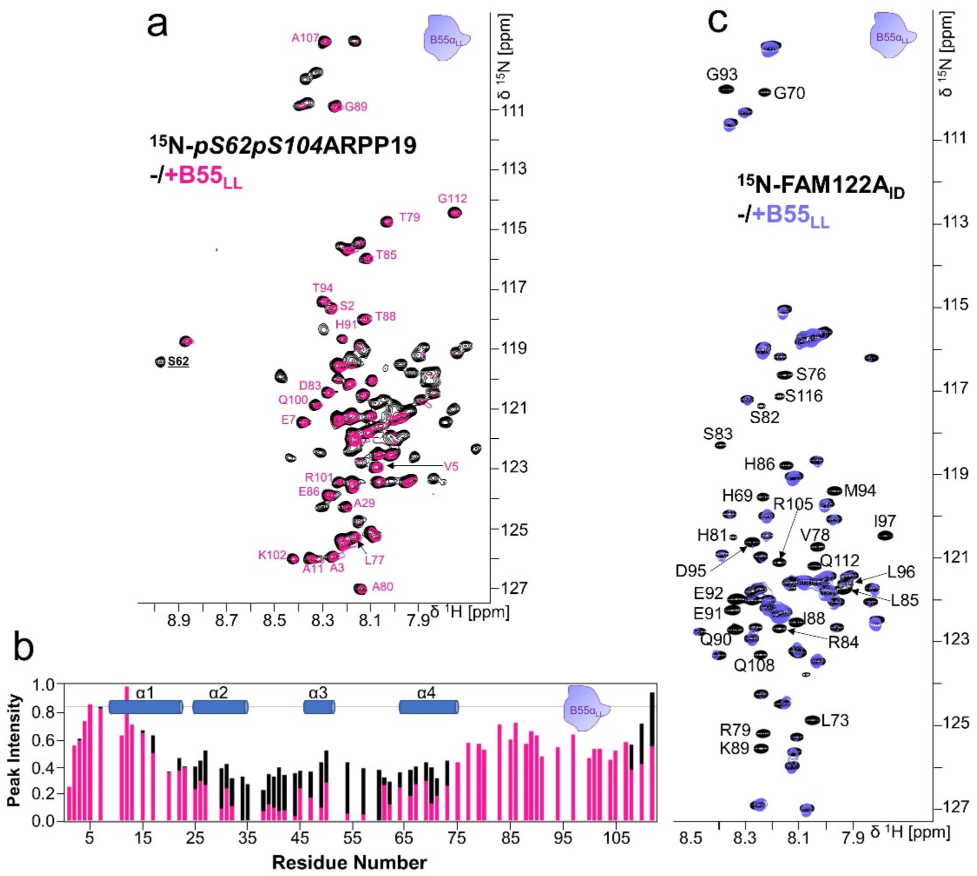

**Extended Data Fig. 6**. **NMR data supporting the interaction of phosphorylated ARPP19 and FAM122A with B55$_{LL}$**. **a**. 2D [$^1$H,$^{15}$N] HSQC spectrum of $^{15}$N-labeled *pS62pS104*ARPP19 alone (black) and in complex with B55$_{LL}$ (pink). pS62 and pS104 labeled for clarity. **b**. Peak intensity vs ARPP19 protein sequence plot for *pS62pS104*ARPP19 alone (black) and when bound to B55$_{LL}$ (pink). Secondary structure elements based on NMR CSI data are indicated. **c**. 2D [$^1$H,$^{15}$N] HSQC spectrum of $^{15}$N-labeled FAM122A$_{ID}$ alone (black) and in complex with B55$_{LL}$ (purple).

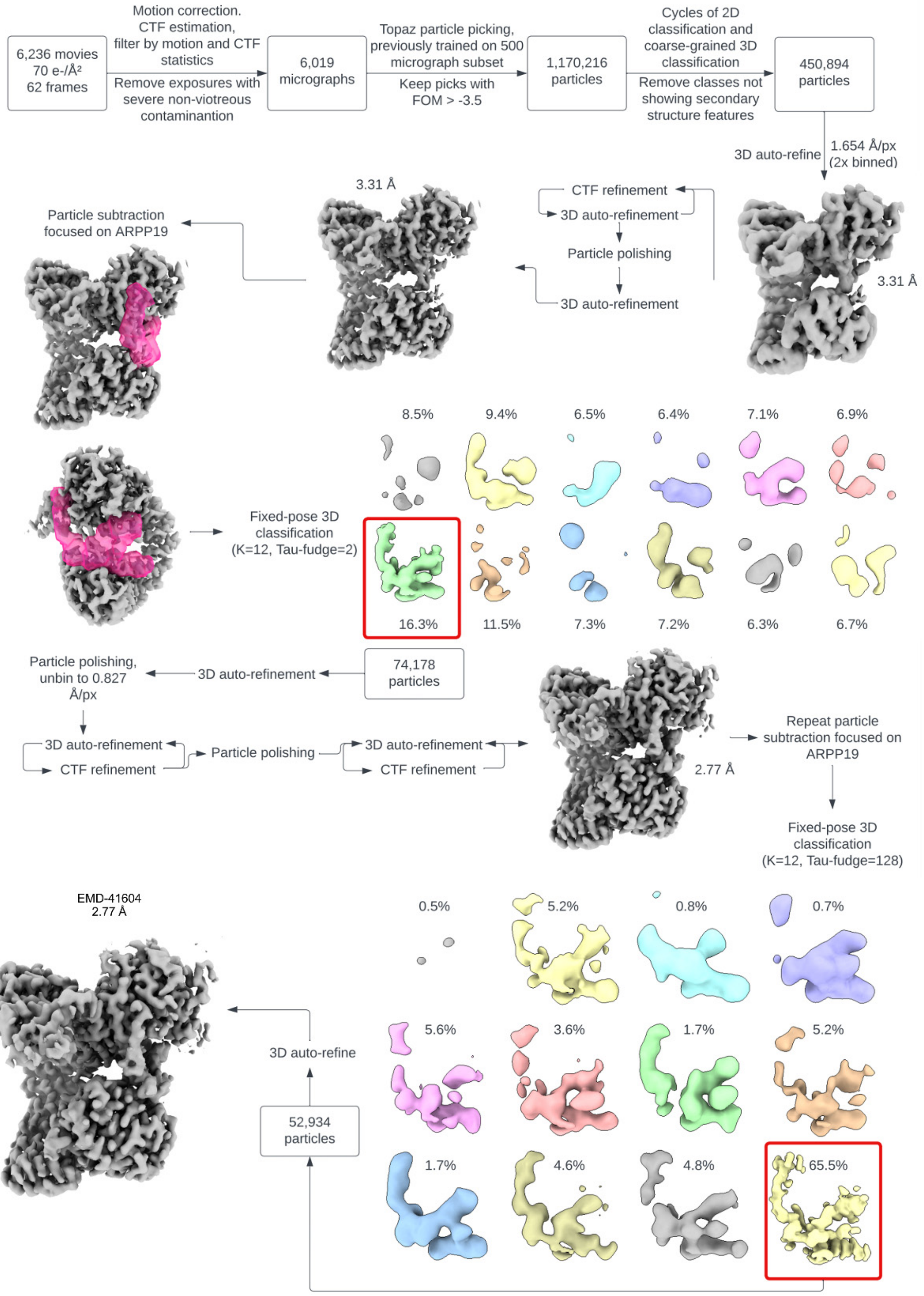

**Extended Data Fig. 7**. **Cryo-EM image processing workflow for PP2A:B55-*tp*ARPP19.** Particle counts and reconstruction resolutions are given at key junctions of the process. "Coarse grained" 3D classification denotes the inclusion of global angular and translational particle pose searches. All resolutions noted are calculated by the gold standard half-maps FSC=0.143 criterion.

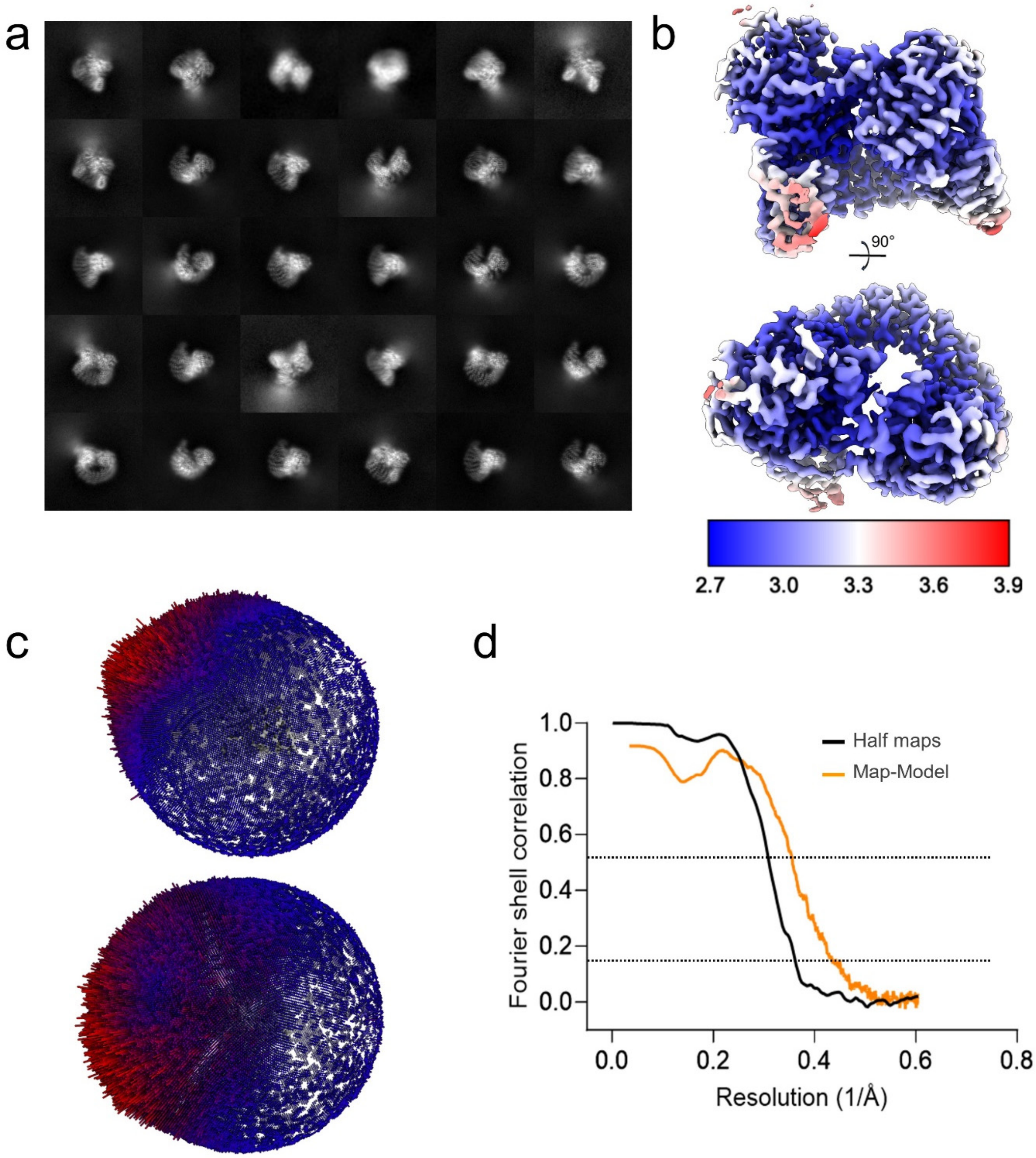

**Extended Data Fig. 8**. **Cryo-EM 2D class averages and maps for PP2A:B55-*tp*ARPP19. a**. Reference-free 2D class averages generated from the 52,934 particles used in the final refinement. **b.** Cryo-EM map, colored by local resolution. **c.** Histograms of the particle pose angular distribution from the input to the final refinement. **d**. The Fourier shell correlation (FSC) for the refinement results. The "gold standard" half-maps FSC was calculated and corrected for masking effects using Relion; the map-model FSC was calculated by PHENIX using a mask around the model based on map resolution. The FSC=0.5 and 0.143 thresholds are marked by dashed lines. For the refinement, the half-maps FSC crosses the 0.143 threshold at 2.77 Å resolution and the map-model FSC crosses the 0.5 threshold at 2.81 Å resolution.

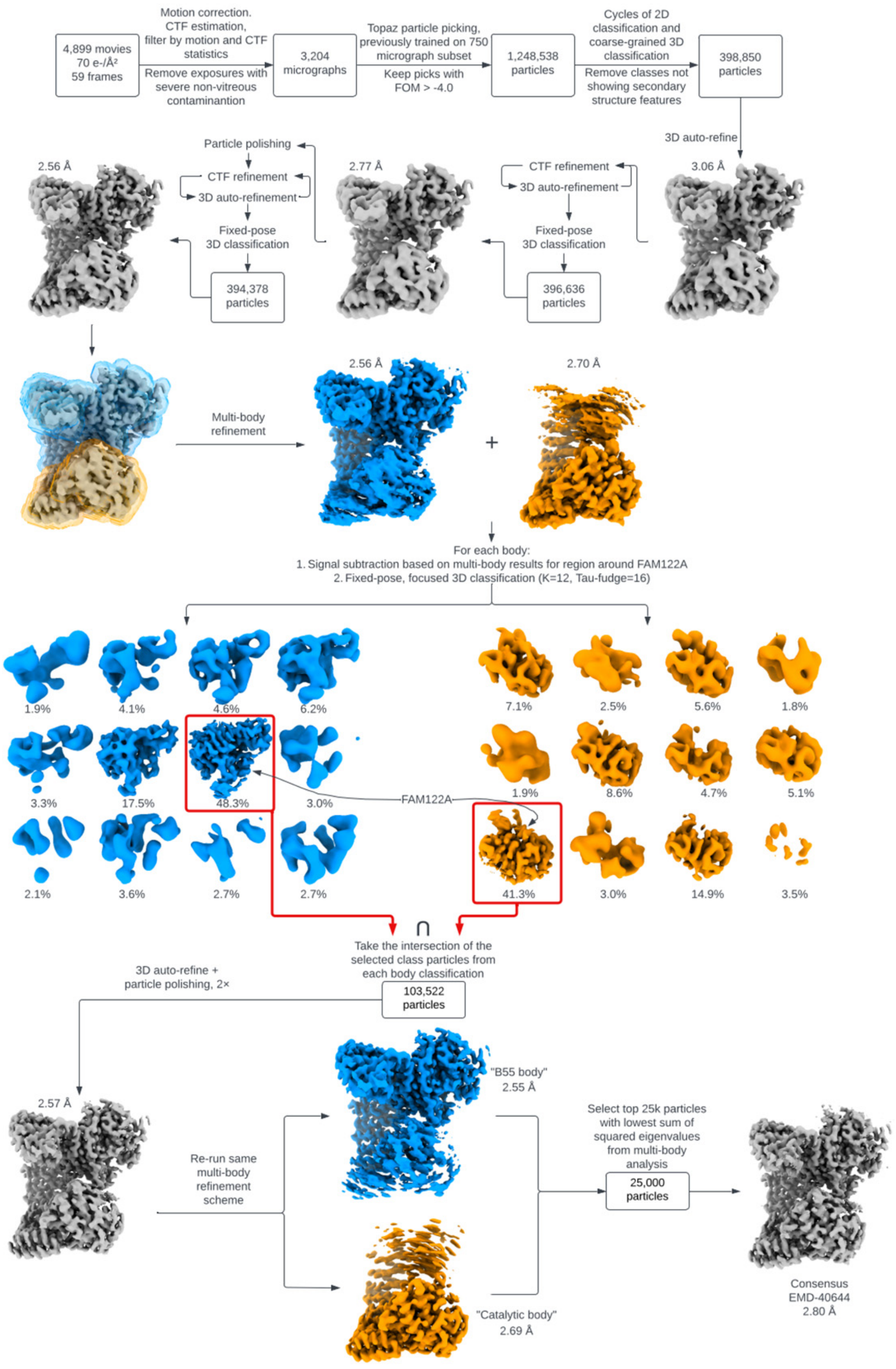

**Extended Data Fig. 9**. **Cryo-EM image processing workflow for PP2A:B55-FAM122A.** Particle counts and reconstruction resolutions are given at key junctions of the process. “Coarse grained” 3D classification denotes the inclusion of global angular and translational particle pose searches. All resolutions noted are calculated by the gold standard half-maps FSC=0.143 criterion.

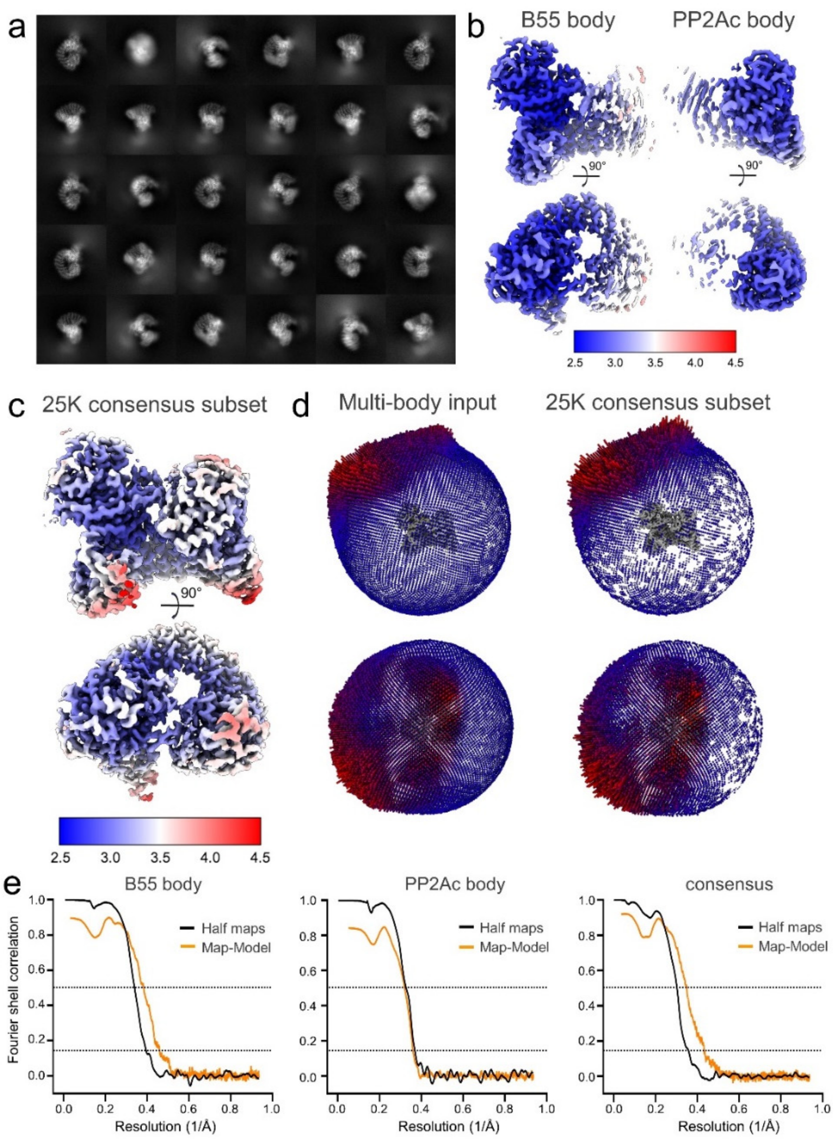

**Extended Data Fig. 10**. **Cryo-EM 2D class averages and maps for PP2A:B55-FAM122A. a**. Reference-free 2D class averages generated from the 103,522 particles used in the final multi-body refinement. **b.** The multi-body refinement results for each body, B55 and PP2Ac, colored by local resolution. **c**. The 25,000-particle subset consensus map, colored by local resolution. **d.** Histograms of the particle pose angular distribution from the input to the final multi-body refinement (left) and the 25,000-particle consensus subset refinement (right). **e**. The Fourier shell correlation (FSC) for the independently refined half-maps (orange) and the full map and atomic model (black) for the multibody refinement results (B55 body, left, PP2Ac body, middle) and the 25,000-particle subset consensus (right). The "gold standard" half-maps FSC was calculated and corrected for masking effects using Relion; the map-model FSC was calculated by PHENIX using a mask around the model based on map resolution. The FSC=0.5 and 0.143 thresholds are marked by dashed lines. For the B55 and PP2Ac multibody refinements, the half-maps FSC crosses the 0.143 threshold at 2.55 Å and 2.69 resolution, respectively, and the map-model FSC crosses the 0.5 threshold at 2.61 Å and 3.12 Å resolution, respectively. For the subset consensus refinement, the half-maps FSC crosses the 0.143 threshold at 2.8 Å resolution and the map-model FSC crosses the 0.5 threshold at 2.86 Å resolution.

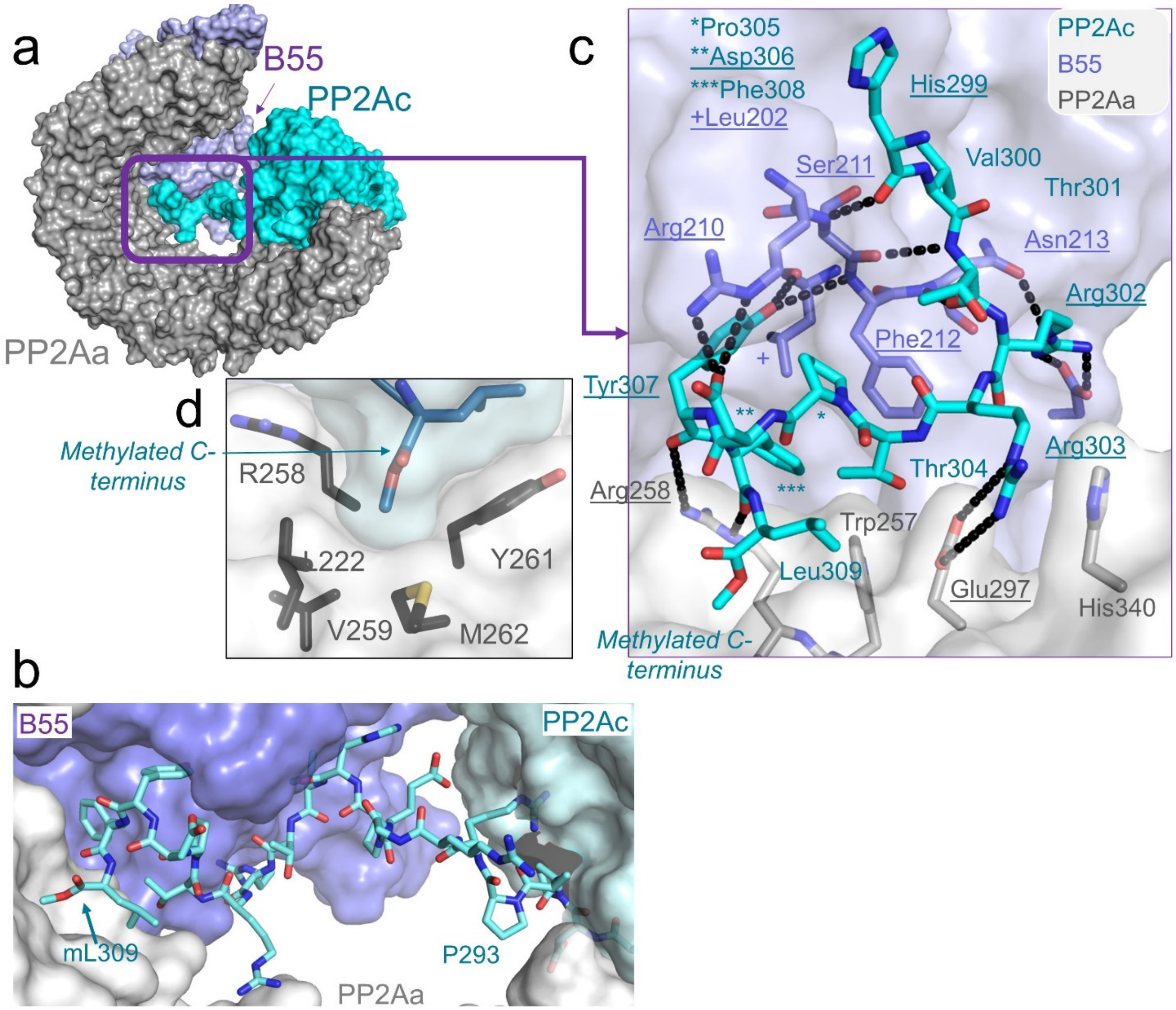

**Extended Data Fig. 11. PP2Ac C-terminus interaction with PP2Aa and B55. a.** The methylated C-terminal tail of PP2Ac (cyan) extends to the opposite side of PP2A (grey) where it interacts at the interface between PP2Aa and B55 (lavender). **b**. Close-up of (a) with the C-terminus shown as sticks and PP2Aa, B55 and the rest of PP2Ac shown as a surface. **c**. Different view of a,b with the PP2Ac C-terminus shown as sticks and PP2Aa and B55 interacting residues also shown as sticks. **d**. Detailed interactions between the methylated C-terminal tail of PP2Ac (cyan, highlighted by purple box in B) with PP2A (grey) and B55 (lavender). Polar/ionic inter-subunit interactions indicated by black dashed lines and the participating residues underlined.

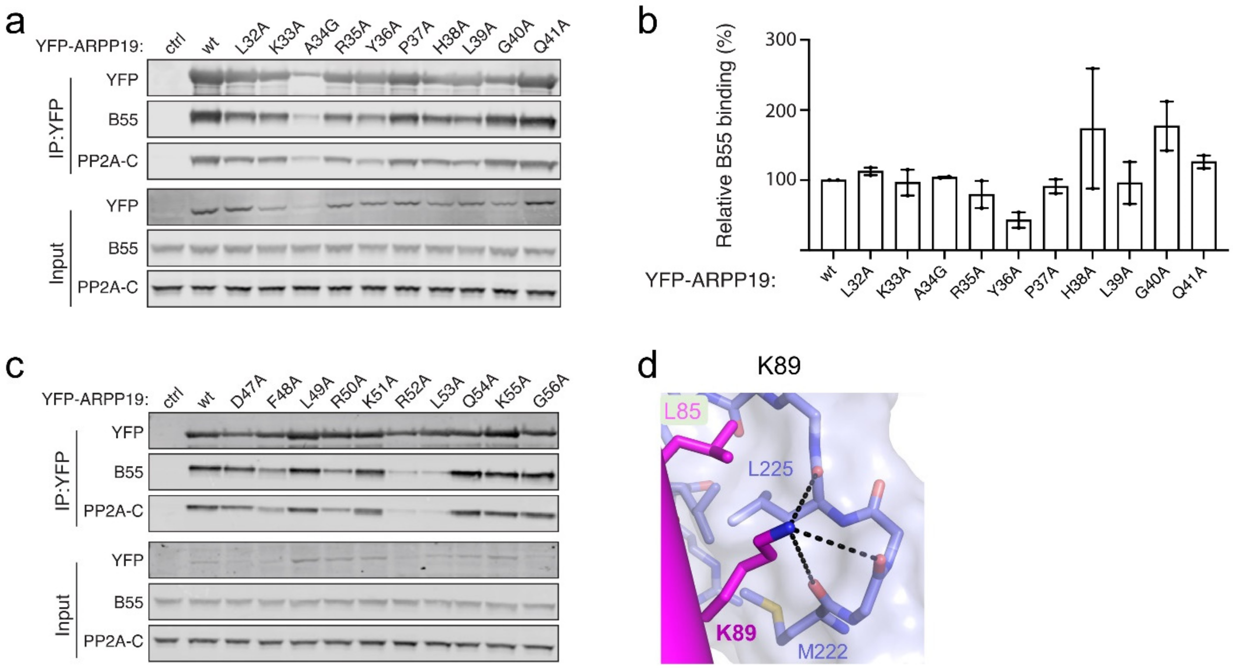

**Extended Data Fig. 12. Pull-down studies of ARPP19 with PP2A and the interaction of FAM122A E92 with B55.** **a.** Alanine scanning mutagenesis of ARPP19 amino acids L32-Q41. The indicated YFP-ARPP19 constructs were transfected into HeLa cells and subsequently immunopurified. Binding efficiency of the YFP-ARPP19 derivatives to B55 and PP2Ac was determined by Western blotting. **b**. Quantification of (a) based on two independent experiments. Error bars are shown as standard error of the mean (SEM). **c**. Alanine scanning mutagenesis of ARPP19 amino acids D47-G56. The indicated YFP-ARPP19 constructs were transfected into HeLa cells and subsequently immunopurified. Binding efficiency of the YFP-ARPP19 derivatives to B55 and PP2Ac was determined by Western blotting. Quantification of (c) based on two independent experiments shown in Fig. 3e. Error bars are shown as standard error of the mean (SEM). **d**. FAM122A K89$_F$ makes polar/ionic interactions (dashes) with multiple residues from B55.

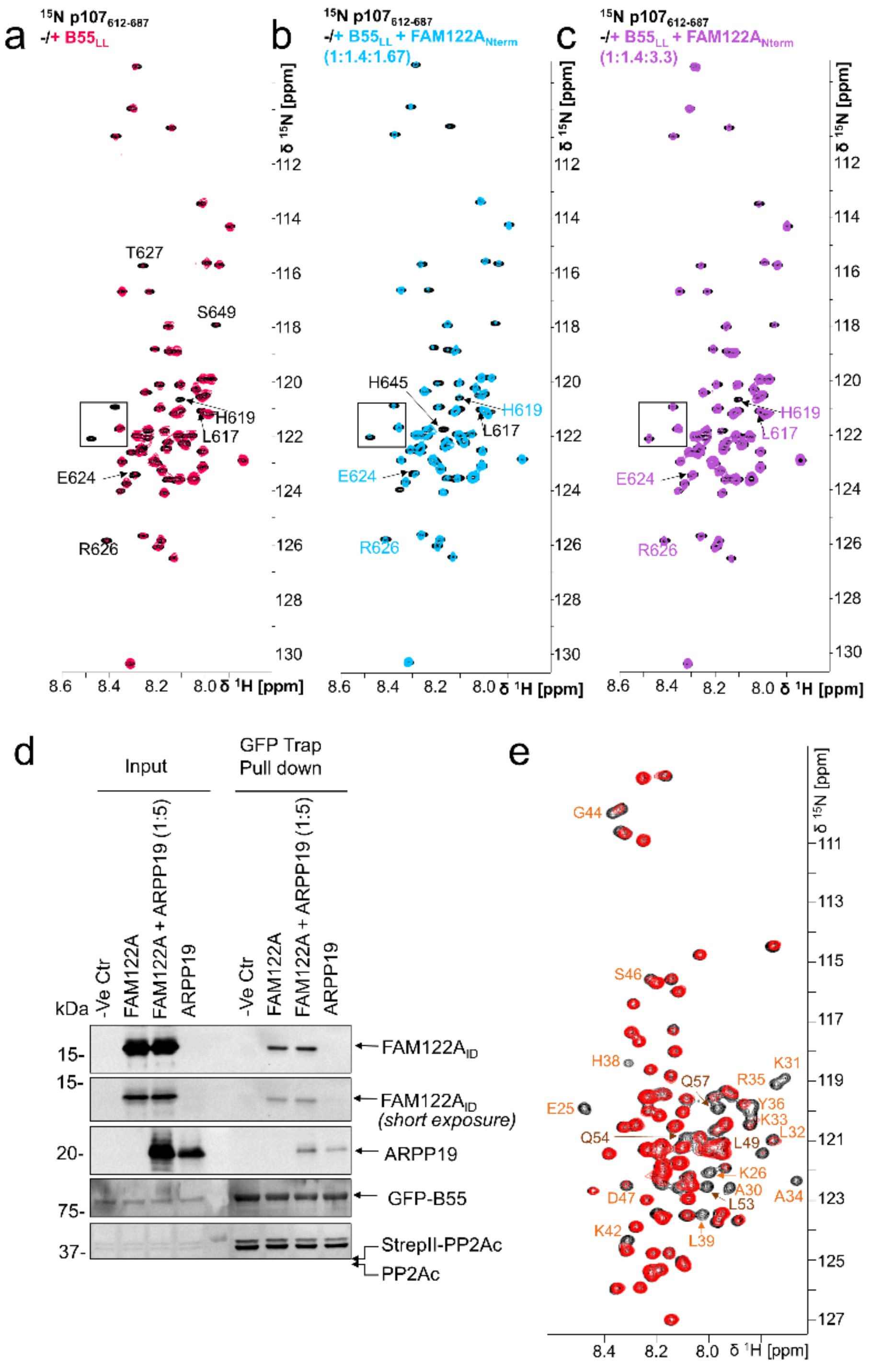

**Extended Data Fig. 13. NMR displacement and pull-down studies show that FAM122A displaces p107 and that FAM122A and *tp*ARPP19 can bind simultaneously. a**. 2D [$^{1}$H,$^{15}$N] HSQC spectrum of $^{15}$N-labeled p107 alone (black) and in complex with B55$_{LL}$ (red) shows that specific p107 residues bind B55$_{LL}$. **b**. The addition of unlabeled FAM122A replaces $^{15}$N-labeled p107 in the identical B55$_{LL}$ binding surface and now starts to appear in the 2D [$^{1}$H,$^{15}$N] HSQC spectrum (light blue) in an identical position as in the 2D [$^{1}$H,$^{15}$N] HSQC spectrum of $^{15}$N-labeled p107 alone (black). **c**. Further addition of unlabeled FAM122A replaces $^{15}$N-labeled p107 in the identical $B55_{LL}$ binding surface and now nearly fully appears in the 2D [$^1H$,$^{15}N$] HSQC spectrum (violet) in an identical position as in the 2D [$^1H$,$^{15}N$] HSQC spectrum of $^{15}N$-labeled p107 alone (black). Black boxes highlight inserts shown in Fig. 3j in the manuscript. **d**. Pulldown assay demonstrating that *tp*S62-ARPP19 does not displace FAM122A when bound to PP2A:B55. Expi293F lysates co-transfected with GFP-B55 and PP2Ac-Strep were incubated with purified PP2Aa and FAM122A alone and with a 5-fold surplus of *tp*S62-ARPP19, pulled-down using a GFP-TRAP and immunoblotted for the indicated proteins. **e**. 2D [$^1H$,$^{15}N$] HSQC spectrum of 4.5 µM $^{15}N$-labeled ARPP19 alone (black) and in complex with 6.9 µM unlabeled $B55_{LL}$ and 22.5 µM unlabeled FAM122A (red) shows that mostly ARPP19 helix α2 $H^N$/N cross peaks stay bound to $B55_{LL}$ (annotated in orange); compare with Fig. 1f without FAM122A.

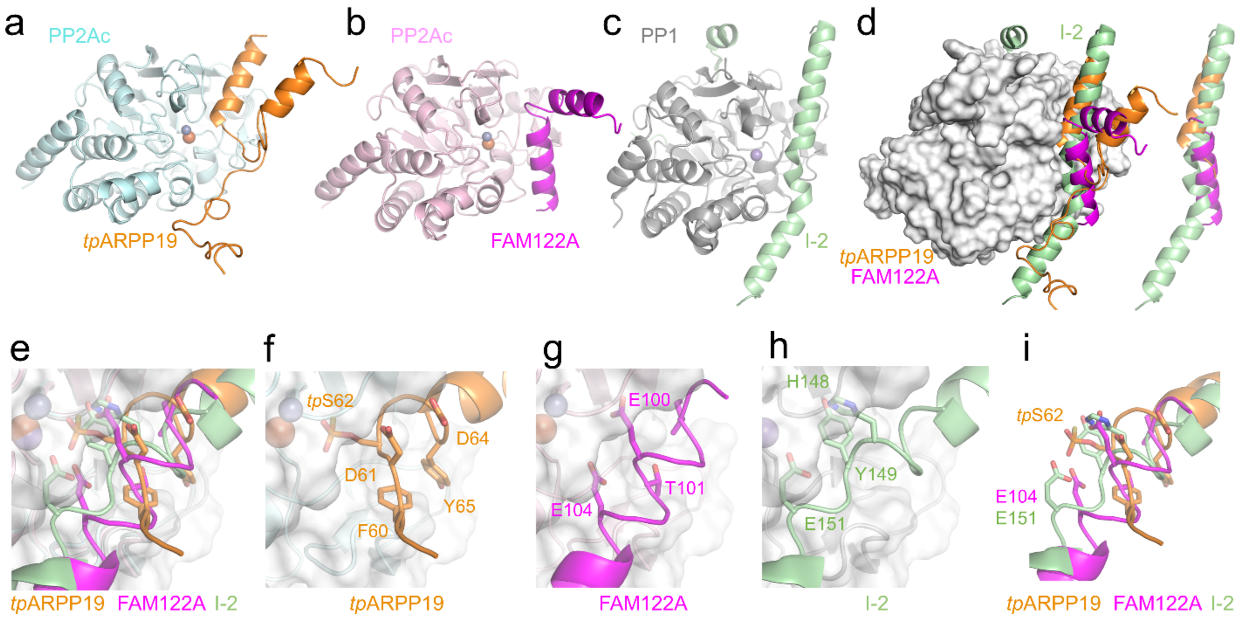

**Extended Data Fig. 14. Overlay of PP2A:B55-*tp*ARPP19, PP2A:B55-FAM122A and PP1:I2.** **a**. *tp*ARPP19 (orange) bound to PP2Ac (cyan). **b**. FAM122A (magenta) bound to PP2Ac (cyan). **c**. I2 (light green) bound to PP1 (grey; pdbid 2OG8). **d**. Overlay of PP2Ac and PP1 bound to *tp*ARPP19, FAM122A and I2, respectively. Colors as in a-c. PP2Ac shown as surface. **e**. Zoom view of (d) with the residues near the active site shown as sticks. Bound metals are shown. **f**. PP2Ac:*tp*ARPP19 in same orientation as e. **g**. PP2Ac:FAM122A in same orientation as e. **h**. PP1:I-2 in same orientation as e. **i**. Same as e but without the PPPs.

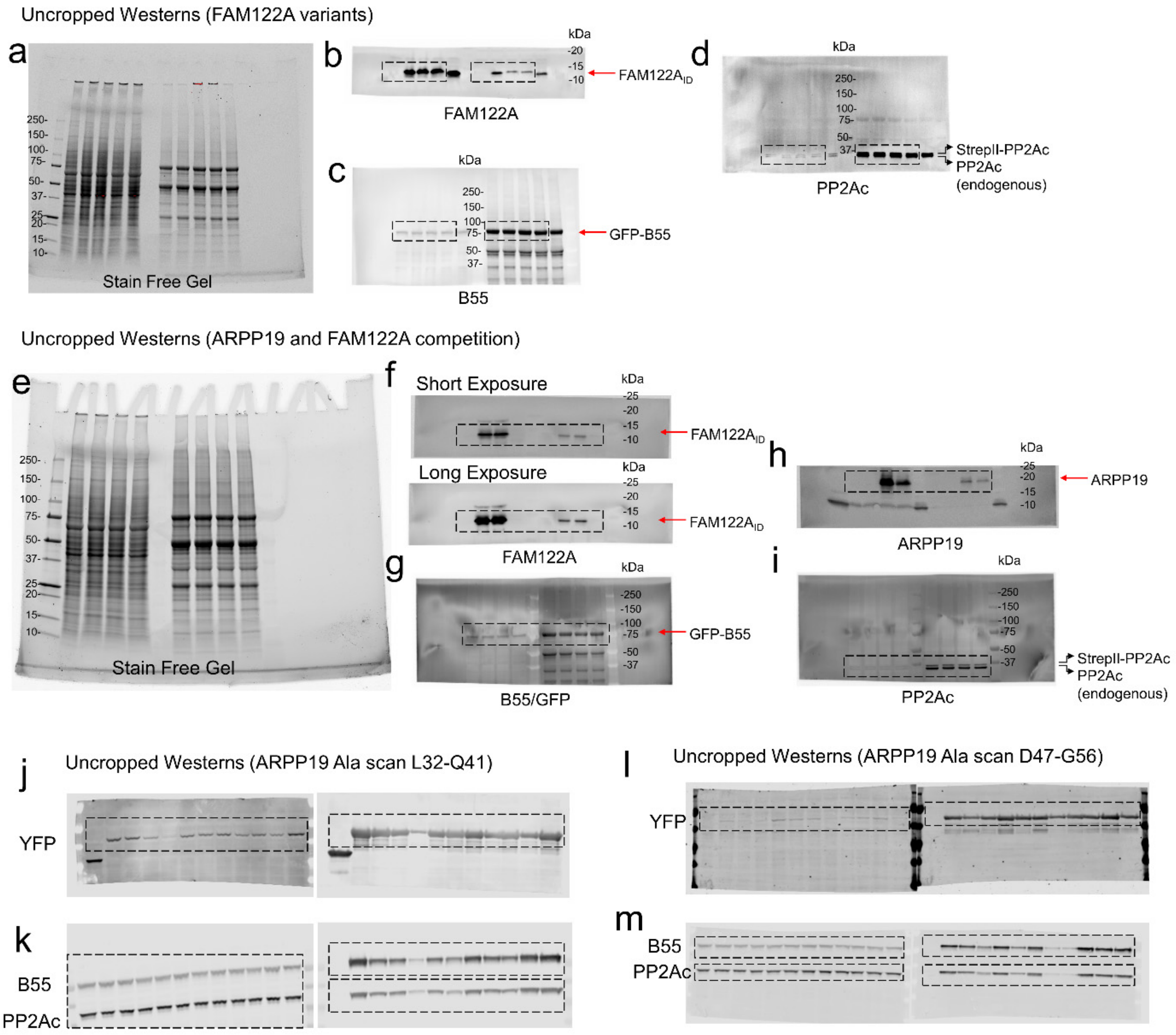

**Extended Data Fig. 15. Uncropped gel and western blot images. a-d.** Refers to Fig 4h. Stain-free SDS-PAGE gel (**a**) used for Western blots (**b-d)**. See legend and panel in 4h for lane identifiers. Bands shown in Fig. 4h indicated by a dashed line box. Molecular weight markers (kDa) shown. **e-i**. Refers to Extended Data Fig. 13d. Stain-free SDS-PAGE (**e**) used for Western blots (**f-i**). See legend and panel in Extended Data Fig. 13d for lane identifiers. Bands shown in Extended Data Fig. 13d indicated by a dashed line box. Molecular weight markers (kDa) shown. **j-m**. Refers to Fig. 3e and Extended Data 12a-c. Uncropped Westerns for ARPP19 Ala scan L32-Q41 **(j,k**) and D47-G56 (**l,m**). Bands shown in Extended Data Fig. 13d indicated by a dashed line box.
